# Within-host evolution of an avirulent *Salmonella* vaccine strain reveals regulatory tuning of virulence functionality without loss of function

**DOI:** 10.64898/2026.11.10.765621

**Authors:** Laëtitia Le Caous, Disha Tandon, Olivier Schären, Luca Beldi, Marco Martinovic, Fereshteh Akbari, Kristýna Filipová, Francisco Osorio-Barrios, Siegfried Hapfelmeier

## Abstract

Non-typhoidal *Salmonella* (NTS) relies on a costly epithelial invasion program encoded in *Salmonella* pathogenicity island 1 (SPI-1) to overcome microbiota-mediated colonization resistance. Because SPI-1 expression is a cooperative trait in which a minority of invasive SPI-1^ON^ cells benefits the wider bacterial population, inflammatory infection models readily select for permanently avirulent “cheater” mutants defective in SPI-1 function. However, while frequently observed during acute infection, isolates from persistent human NTS infection rarely exhibit obvious SPI-1 loss-of-function mutations, raising the question of whether SPI-1 contributes to bacterial fitness beyond induction of acute intestinal inflammation and tissue persistence. Here, we addressed this question using a metabolically attenuated yet epithelial-invasive auxotrophic *Salmonella* serovar Typhimurium (STm) vaccine strain (STm^Aux^) that persistently colonizes the murine intestine without causing overt inflammatory pathology or tissue infection while retaining SPI-1-dependent host interaction and immunogenicity. Following within-host evolution for up to five years, STm^Aux^ populations did not accumulate classical *hilD* or other obvious SPI-1 loss-of-function cheater mutants. Instead, parallel evolution repeatedly targeted the BarA/SirA two-component regulatory system, closely mirroring previous observations in persistent human NTS infections. Functional analyses demonstrated that evolved *barA* and *sirA* alleles tune rather than abolish SPI-1 function by sharply reducing the proportion of SPI-1^ON^ cells while preserving their individual epithelial invasion competence. Reconstructed in a prototrophic strain background, early evolved isolates retained full intestinal and systemic virulence *in vivo* despite exhibiting strongly reduced SPI-1^ON^ frequencies *in vitro*. Remarkably, identical *sirA* mutant alleles also emerged in a *hilD-*deficient strain background, indicating that selection for these adaptive variants is not contingent on canonical HilD-dependent SPI-1 activation. During prolonged evolution, additional mutations accumulated that were associated with loss of systemic virulence despite preservation of epithelial invasiveness, suggesting subsequent adaptation affecting host-interaction pathways beyond SPI-1-mediated virulence. Together, these findings identify regulatory tuning of the BarA/SirA control network as a conserved evolutionary trajectory during prolonged NTS persistence. Rather than favoring classical cheater-driven loss of virulence functions, long-term adaptation preserves substantial SPI-1-dependent host interaction while reducing the proportion of phenotypically virulent SPI-1^ON^ cells.

## Introduction

Intestinal mucosal virulence of non-typhoidal *Salmonella* enterica (NTS) relies on the type 3 secretion system encoded by *Salmonella* pathogenicity island 1 (SPI-1), which enables epithelial cell invasion and initiation of inflammatory host-pathogen interactions in the intestinal mucosa [1,3]. SPI-1 expression is a classical cooperative trait. A dedicated phenotypically virulent SPI-1-expressing (SPI-1^ON^) subpopulation engages in the metabolically costly virulence program required for epithelial cell invasion. These invasive but slow-growing cells actively cross the epithelial barrier during early-stage infection, where they are engulfed by host phagocytes and largely eliminated, constituting a genuinely self-sacrificial subpopulation. In contrast, the remaining fast-growing phenotypically avirulent SPI-1^OFF^ population in the intestinal lumen benefits from inflammation-derived nutrients generated as a consequence of SPI-1-dependent host interaction, thereby exploiting a public good produced by the invasive subpopulation [1-3]. This division of labor inherently exposes NTS populations to cheating, as spontaneous permanently avirulent “cheater” mutants can potentially benefit from inflammation without contributing to its induction [1,3,4].

Despite this vulnerability, complete loss of SPI-1 function is rarely observed during long-term persistence. Experimental and clinical studies indicate that SPI-1-deficient “cheater” mutants may transiently expand during acute infection but are frequently outcompeted as the intestinal ecosystem recovers [3,4,5]. Multiple mechanisms have been proposed to contribute to the long-term stabilization of cooperative virulence, including competition from bistable wild-type SPI-1^OFF^ populations that maintain a phenotypically virulent SPI-1^ON^ subpopulation, reseeding SPI-1 proficient bacteria from tissue reservoirs, and microbiota competition-mediated selection favoring maintenance of SPI-1 function [5,6]. These observations suggest that while SPI-1 expression is evolutionarily dynamic, complete loss of virulence function is strongly counter-selected during prolonged host colonization and host-host transmission [5,7].

This raises the possibility that long-term intestinal persistence may favor regulatory optimization of virulence expression rather than simple loss of virulence functions, particularly under chronic or asymptomatic infection conditions where overt inflammatory pathology is absent. Consistent with this idea, clinical studies of prolonged persistent and recurrent NTS infection in humans have not identified widespread accumulation of classical SPI-1-deficient cheater genotypes such as loss-of-function mutations in the SPI-1 master regulator gene *hilD* that are commonly observed during acute disease [1,5,7]. Instead, prolonged carriage is frequently associated with mutations in global regulatory systems, most prominently the BarA/SirA two-component regulatory pathway [8]. These persistence-associated variants were reported to reduce SPI-1 gene expression, yet their physiological significance and precise impact on SPI-1-dependent functions remain poorly understood. Whether such mutations represent attenuated loss-of-function (“cheater”-like) variants or instead mediate more subtle tuning of virulence expression is currently unknown.

This question has important implications not only for understanding pathogen evolution during infection but also for the development of live-attenuated oral *Salmonella* vaccines. In previous work, we described an auxotrophic *Salmonella* serovar Typhimurium strain (STm^Aux^) engineered to be strictly dependent on the peptidoglycan precursors D-alanine (D-Ala) and *meso*-diaminopimelic acid (*m-*DAP) [9]. Although fully attenuated and unable to cause detectable intestinal inflammation or systemic infection, STm^Aux^ retains SPI-1-dependent epithelial invasiveness and host-interaction capacity. Remarkably, STm^Aux^ elicits high-avidity mucosal IgA responses and robust protective immunity in a manner that remains dependent on SPI-1 function despite the complete absence of inflammatory pathology [9]. Local metabolic cross-feeding by the intestinal microbiota supports luminal colonization of STm^Aux^ while remaining insufficient to rescue invasive tissue infection, thereby uncoupling SPI-1-dependent host interaction and immunogenicity from virulence [9]. Our previous study further demonstrated stable maintenance of the engineered auxotrophic attenuation during prolonged intestinal colonization [9]. Whether the SPI-1-dependent host interaction remains evolutionarily advantageous in the absence of virulence is unknown.

These properties make STm^Aux^ a unique and relevant model to investigate how a pathogen-mimetic, but inflammation-silent *Salmonella* strain evolves during long-term residence in the gut. Specifically, does avirulence render SPI-1 function evolutionarily dispensable, favoring the emergence of classical avirulent cheater genotypes, or does SPI-1-dependent host interaction continue to confer a selective advantage despite the absence of overt pathology?

To address this question, we investigated long-term within-host evolution of STm^Aux^ during persistent intestinal colonization of gnotobiotic mice harboring the defined Oligo-MM^12^ microbiota [10]. Rather than accumulating obvious SPI-1 cheater genotypes such as *hilD* loss-of-function mutations, evolving STm^Aux^ populations repeatedly acquired highly parallel mutations in the BarA/SirA regulatory system. Remarkably, several of these variants resembled mutations previously identified in persistent human NTS infections [8]. Functional analyses revealed that adaptive BarA/SirA variants quantitatively remodel SPI-1 bistability, markedly reducing the fraction of SPI-1-expressing cells while preserving a functional invasion-competent subpopulation. Together, these findings identify regulatory tuning of a major virulence control network as a conserved evolutionary trajectory during prolonged NTS persistence.

## Results

### Gnotobiotic Oligo-MM^12^ mice enable long-term STm^Aux^ colonization and within-host evolution

To investigate within-host evolution of STm^Aux^ we conducted long-term colonization experiments in gnotobiotic Oligo-MM^12^ mice harboring a 12-species commensal microbiota with low colonization resistance against STm. This model enables intestinal colonization without antibiotic pre-treatment [10] and, through local metabolic cross-feeding by the microbiota [9], supports sustained gut luminal colonization of STm^Aux^ despite its strict auxotrophy. Adult male and female mice were individually inoculated with 10⁷ colony-forming units (CFU) of STm^Aux^ and, following establishment of stable intestinal colonization for 4 weeks, were bred to initiate serial vertical transmission lineages (founder generation F0; **Fig. 1A**).

**Figure 1.**
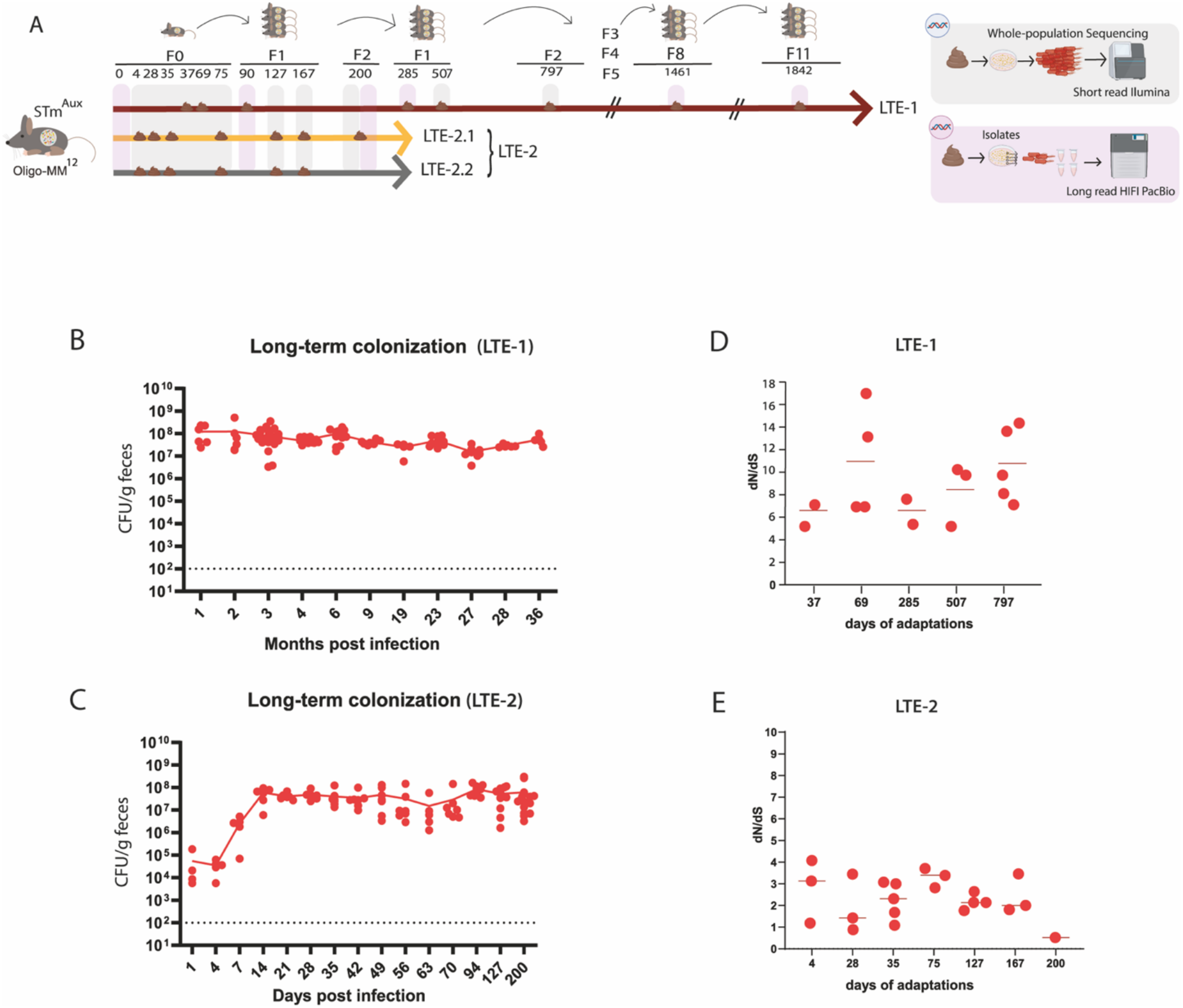
Prolonged intestinal colonization reveals adaptive genomic evolution of STm^Aux^. **(A)** Experimental design of the long-term evolution (LTE) studies. Three independent STm^Aux^ lineages were evolved during prolonged intestinal colonization of Oligo-MM^12^ mice: LTE-1 (brown) across 12 host generations (F0-F11) and LTE-2.1 (orange) and LTE-2.2 (grey) across three host generations (F0-F2). Fecal samples were collected longitudinally and analyzed either by population-level Illumina sequencing (grey) or by PacBio HiFi sequencing of individual re-isolated strains (pink). Four evolved STm^Aux^ re-isolates (90d, 200d, 1461d, and 1842d STm^Aux^) carrying representative adaptive mutations were selected for downstream characterization. **(B)** Fecal bacterial loads of LTE-1 quantified by CFU plating on selective media and shown across months post-infection. **(C)** Fecal bacterial loads of LTE-2 quantified similarly and shown across days post-infection. (**D-E**) Ratio of non-synonymous to synonymous mutations (dN/dS) in LTE-1 (D) and LTE-2 (E) populations. Each symbol represents a single mouse-derived population. Horizontal bars indicate means. The dotted lines indicate the detection limit.

A first long-term evolution experiment (LTE-1; **Fig. 1A**) spanned approximately five years (1842 days) of continuous colonization across 12 host generations (founder generation F0 and 11 filial generations F1-11) (**Supp. Fig. 1**). A second experiment (LTE-2) extended over 200 days and three host generations and, through experimental cage separation, generated two independent evolutionary STm^Aux^ lineages (LTE-2.1 and LTE-2.2; **Fig. 1A**).

Fecal STm^Aux^ populations were cultivated on selective agar to quantify colonization densities (**Fig. 1B, C**) and to isolate STm^Aux^ bacterial DNA for short-read Illumina metagenome sequencing to study mutant allele frequency dynamics within evolving STm^Aux^ populations. Complementing these population-level genome analyses, evolved STm^Aux^ clones were re-isolated at four time points (90, 200, 1461, and 1842 days; **Fig.1A**) and subjected to long-read PacBio HiFi genome sequencing, enabling high-resolution strain-level and functional genetic analysis. Together, this combined experimental framework enabled reconstruction of the temporal dynamics of STm^Aux^ within-host evolution across independent evolutionary lineages and linked population-level allele dynamics to defined evolved genotypes and strains.

### Selection-driven, adaptive within-host evolution of STm^Aux^

We first sought to establish whether the genetic diversification of STm^Aux^ during long-term intestinal colonization was consistent with selection-driven adaptive evolution. A key signature of such selection is an excess of non-synonymous over synonymous mutations, as amino acid-changing substitutions are more likely to be retained when they confer a fitness advantage, whereas synonymous substitutions are often used as a proxy for mutations with weaker effects on protein function [11,12]. Consistent with this expectation, both LTE-1 and LTE-2 exhibited a non-synonymous-to-synonymous mutation ratio greater than one (**Fig. 1D, E**), supporting the hypothesis that bacterial evolution in both experiments was selection-driven rather than neutral [11-13]. To determine whether this molecular signature was accompanied by a measurable fitness benefit *in vivo*, we compared a late-evolved STm^Aux^ re-isolate with its ancestor in a competitive intestinal colonization experiment. A day-1461 re-isolate from LTE-1 rapidly outcompeted the ancestral STm^Aux^ strain (**Fig. 2**), providing direct evidence that within-host evolution had led to an increased *in vivo* fitness phenotype. These data are consistent with adaptive evolution of STm^Aux^ during long-term colonization.

**Figure 2.**
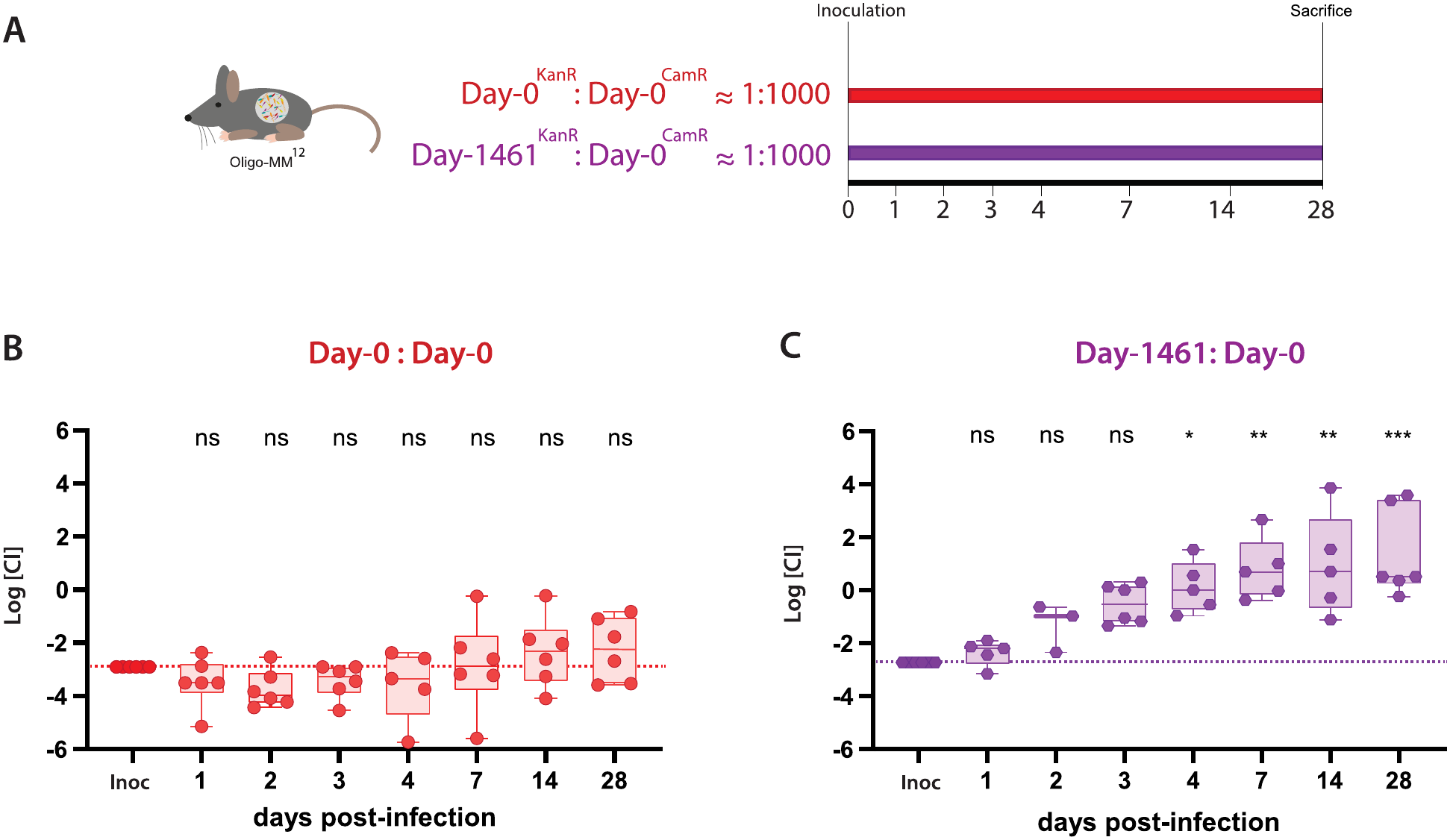
Long term–evolved STm^Aux^ strains exhibit increased fitness in the Oligo-MM^12^ mouse model. **(A)** Schematic of the competitive infection experiment. Oligo-MM^12^ mice were gavaged with a total of 10⁸ CFU of either a mix of Day-0^KanR^ : Day-0^CamR^ (non-adapted STm^Aux^ vs non-adapted STm^Aux^) at a ratio of ∼1:1000, serving as negative control, or Day-1461d^KanR^: Day-0^CamR^ (1461d STm^Aux^ vs non-adapted STm^Aux^) at a ratio of ∼ 1:1000, n=6 per group. Fresh fecal samples were collected longitudinally at the indicated time points. Mice were sacrificed at day 28 post-inoculation, and bacterial loads were quantified by plating on selective media containing kanamycin or chloramphenicol to distinguish competing strains. **(B)** Competitive index (CI) was calculated as the ratio of Day-0^KanR^/Day-0^CamR^ for each day post inoculation. **(C)** Competitive index (CI) was calculated as the ratio of Day-1461^KanR^/Day-0^CamR^ for each day post inoculation. At some time points n <6 mice are shown, because not all mice produced fecal samples at all time points. Inoc, inoculum; dotted lines indicate the competitive index in the bacterial inoculum. Data were analyzed using the Kruskal–Wallis test followed by Dunn’s multiple-comparisons test. p values: ns p > 0.05; *p < 0.05; **p < 0.01; ***p < 0.001.

### Parallel mutations targeting the BarA/SirA two-component system and convergence across independent studies

The experimental evolution of enterobacterial populations *in vivo*, like *in vitro*, is characterized by soft selective sweeps in which multiple adaptive lineages coexist, rather than rapid fixation of a single dominant clone [13-15]. Genomic sequencing consequently revealed an extensive diversity of low-frequency mutant alleles across host individuals, sampling time points and independent long-term evolution experiments.

Within this genomic diversification, STm^Aux^ evolution repeatedly targeted genes encoding components of regulatory systems involved in controlling virulence and cellular physiology in *Salmonella* [16-18]. Among the most recurrent variants, *gatR* mutations reached high frequencies in LTE-1 and LTE-2 [19], while mutations in the *sirA/barA* regulatory system became selected early during colonization in both experiments [20,21]. Mutations in *envZ/ompR* additionally highlighted the possible implication of environmental sensing through global two-component regulatory systems, which are known to link environmental adaptation and virulence gene regulation [22,23].

Our findings in part paralleled previous studies studying STm within-host evolution in mice. In one study using the streptomycin-pretreated mouse model, the glycerol-3-phosphate transporter gene *glpT* was found mutated in approximately 70% of evolved STm isolates [5,24]. Similarly, in our study, 100% of LTE-1 re-isolates studied at day 1842 carried a large deletion encompassing *glpA*-*glpT*-*glpQ* (**Supplementary Tables 1 and 2**). We also reproduced the occurrence of 7-bp frameshift variants in *oafA*, previously reported by Emma Slack and colleagues as an intestinal IgA escape LPS O-Antigen phase variant transiently selected during enteric STm infection [25].

Several previous genomic studies of NTS in murine enteric infection models and clinical isolates established mutations in the SPI-1 master regulator *hilD* and the two-component system *sirA/barA* as particularly frequent targets for “cheater” mutations that abolish SPI-1-mediated mucosal virulence and thereby provide a temporary adaptive benefit within the inflamed intestine [1,5,7]. However, against our initial hypothesis that avirulent STm^Aux^ would also strongly benefit from abolition of costly expression of SPI-1, mutations in *hilD* were not predominant during within-host evolution of STm^Aux^. Moreover, while mutations in *sirA/barA* did recurrently accumulate in evolving STm^Aux^, these were primarily conservative indel or amino acid substitutions rather than frameshift or nonsense mutations, suggesting they were not simply loss-of-function mutations. In this point our results differed from previous findings in acute mouse models but showed striking parallels with recent clinical study of within-host evolution during long-term persistent human NTS infection [8].

In LTE-1, a conservative 6-bp in-frame deletion in *sirA*, *sirA^c^*^.548_553del^ (p.184_185del; hereafter SirA^Δ184-185^) arose early during colonization and rapidly reached fixation (**Fig. 3C, D**). The deleted residues map to the C-terminal helix–turn–helix DNA-binding domain of SirA, a region implicated in promoter recognition and transcriptional control (**Fig. 3C**) [21,31,32].

**Figure 3.**
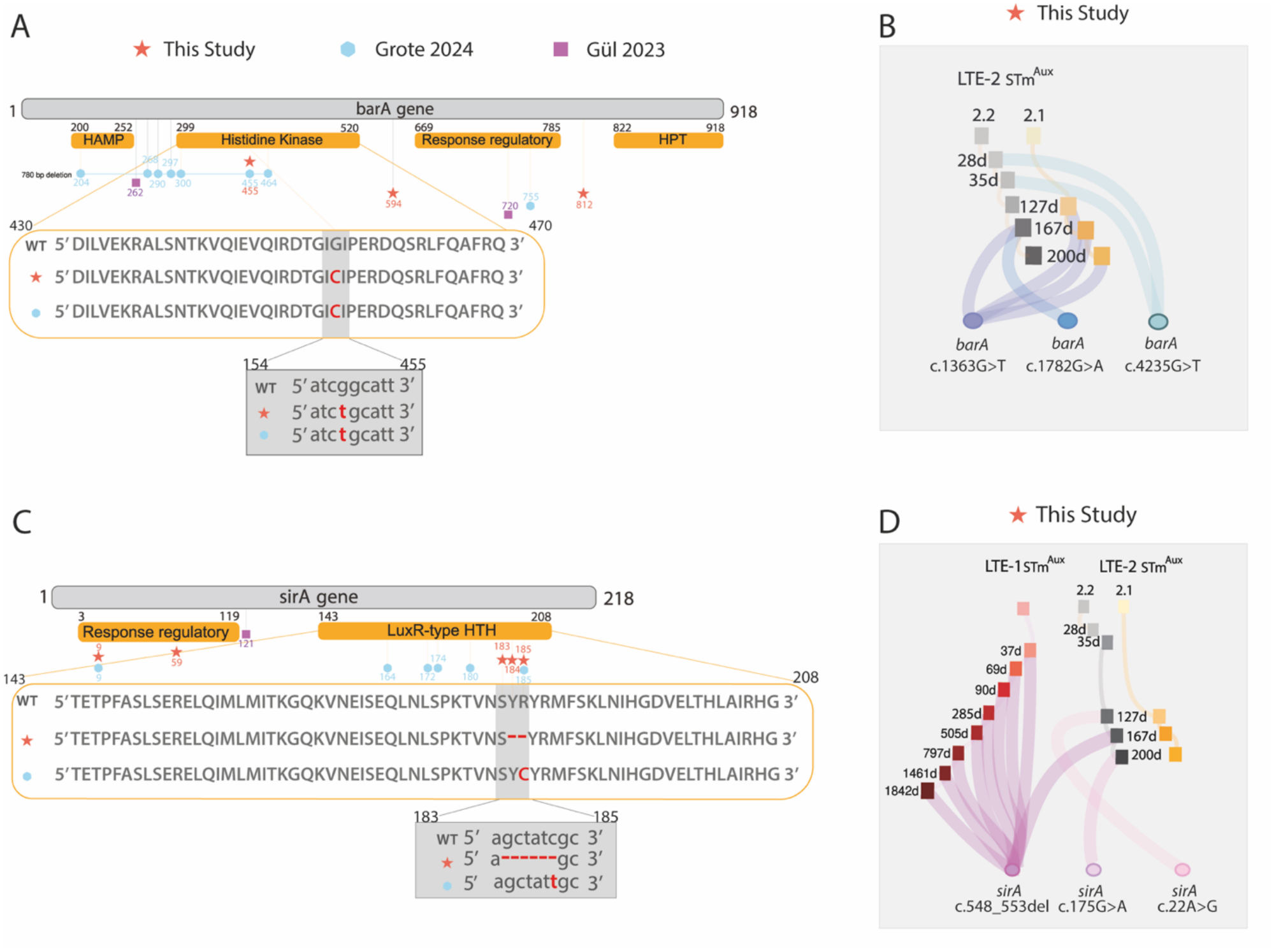
Parallel selection of *barA* and *sirA* during long-term *in vivo* adaptation of non-typhoidal *Salmonella*. **(A)**, **(C)** Schematic representation of the *barA* (A) and *sirA* (C) loci showing the major functional domains annotated from Interpro [31] (highlighted in orange). Mutations identified in this study are indicated with red stars, while those reported in Gül et al., 2023 are shown as purple squares, and mutations from Grote et al., 2024 are depicted as blue circles. Mutations are labelled according to the position of the affected amino acid in the protein sequence. Related *barA* and *sirA* mutations that were identified in the present and Grote et al. study are additionally depicted as amino acid (orange bars) and nucleotide (grey boxes) sequence alignments. **(B)**, **(D)** Sankey diagrams illustrating the emergence and temporal dynamics of *barA* (B) and *sirA* (D) mutations across the three long-term evolution experiments: LTE-1 (red), LTE-2.1 (orange) and LTE-2.2 (grey). Mutations in *barA* are shown in different shades of blue, with each shade representing a distinct variant linked to one of the three LTE lineages and the day on which the mutation occurred. The same scheme applies to *sirA* with different shades of pink. Sankey diagrams were generated using sankeymatic.com.

In LTE-2.2, the identical *sirA*^c.548-553del^ allele emerged independently, although only transiently and at intermediate frequency (≈ 17 %) in one mouse by day 167 before being displaced by an alternative *sirA* missense mutant allele affecting the N-terminal receiver domain (*sirA*^c.175G>A^; p.Gly59Ser) (**Fig. 3C, D**). Earlier during evolution of the same lineage, yet another receiver domain substitution, *sirA*^c.22A>G^ (p.Asp9Gly) transiently emerged (**Fig. 3C, D**).

Parallel to recurrent *sirA* evolution in LTE-2.2, multiple independent *barA* alleles emerged across LTE-2.2 (**Fig. 3A, B).** In contrast to LTE-1 and LTE-2.2, LTE-2.1 was dominated by a single *barA* missense mutation, *barA*^c.1363G>T^ (p.Gly455Cys, hereafter BarA^Gly455Cys^), which reached high frequency by the experimental endpoint and was confirmed in multiple independently re-isolated clones by whole-genome sequencing (**Fig. 3A, B**). Importantly, the spectrum of adaptive *sirA* and *barA* mutations was dominated by conservative amino-acid substitutions or small in-frame deletions rather than nonsense mutations or disruptive frameshifts. Together, these findings identify adaptive remodeling of the BarA/SirA regulatory system as a major evolutionary trajectory during long-term STm^Aux^ colonization rather than complete loss of virulence function.

The mutant alleles or *sirA/barA* that recurrently evolved during long-term colonization of STm^Aux^ in our mouse experiments displayed striking convergence with long-term within-host evolution of natural NTS in long-term persistently infected humans [8], not only at the level of the genes, but also at the protein domains targeted and sometimes even resulted in identical or very similar amino acid changes.

Most notably, the BarA^Gly455Cys^ substitution that predominated in LTE-2.1 was also reported in a patient persistently infected by non-typhoidal *Salmonella* (NTS) serovar Muenchen [8]. Convergence was also apparent in SirA: one within-host evolved mutation identified in the human cohort was an amino acid substitution of Arg185 within the C-terminal DNA-binding domain, one of the two amino acids deleted by the conservative SirA^Δ184–185^ deletion that reached fixation in STm^Aux^ LTE-1 and transiently emerged during LTE-2.2 (**Fig. 3C**). A second *sirA* mutant allele that emerged in both studies was the SirA^Asp9Gly^ receiver domain substitution that we observed in STm^Aux^ LTE2.2.

Together, these findings reveal convergent evolution of the BarA/SirA regulatory system across experimental and clinical settings, encompassing both recurrent amino-acid substitutions and independent mutations targeting the same functional domain. This convergence is consistent with shared selective constraints on regulatory pathways during prolonged intestinal colonization, while leaving open whether the underlying mutations confer similar phenotypic effects in each genetic and host context.

### Evolved *barA/sirA* mutations tune SPI-1 expression while preserving cell-invasion competence

The BarA/SirA two-component system is a central upstream regulator of the HilD regulon controlling SPI-1 and associated virulence gene expression and has repeatedly been implicated in virulence attenuation during within-host evolution of NTS [5, 7,8, 21, 28]. We therefore asked whether the recurrent *sirA* and *barA* mutations selected during long-term STm^Aux^ colonization modulate SPI-1– dependent host cell interaction and virulence-associated functions.

To enable functional analysis of evolved genotypes, we applied an ex-auxotroph reverse-engineering strategy in which the auxotrophic deletions in *asd* and *dadX*/*alr* were restored to wild-type configuration in selected evolved STm^Aux^ strains as well as in the ancestral STm^Aux^ strain background and corresponding isogenic mutant controls, generating replication-competent “STm^Ex-Aux”^ derivatives. This approach allows assessment of SPI-1-dependent cell invasion and *in vivo* virulence phenotypes independently of the metabolic attenuation caused by auxotrophy.

SPI-1-dependent epithelial cell invasion was quantified using gentamicin protection assays in HeLa cells. SPI-1 T3SS-competent wild-type STm, STm^Aux^, and STm^Ex-Aux^ served as invasive positive controls, whereas SPI-1 T3SS-deficient STm^Aux^ ^Δ*invC*^ and STm^Ex-Aux^ ^Δ*invC*^ mutants were included as fully non-invasive negative controls. Multiple evolved STm^Aux^ re-isolates were analyzed and grouped according to their *sirA* and *barA* genotype (**Fig. 4A**). To distinguish altered regulatory activity from complete loss of BarA/SirA function, isogenic Δ*barA* and Δ*sirA* deletion mutants constructed in the ancestral STm^Aux^ strain background were included. In addition, the specific contribution of the evolved alleles was assessed by reconstructing either *sirA*^c.548_553del^ (SirA^Δ184–185^) or *barA*^c.1363G>T^ (BarA^Gly455Cys^) individually in the ancestral STm^Aux^ strain background (**Fig. 4A**).

**Figure 4.**
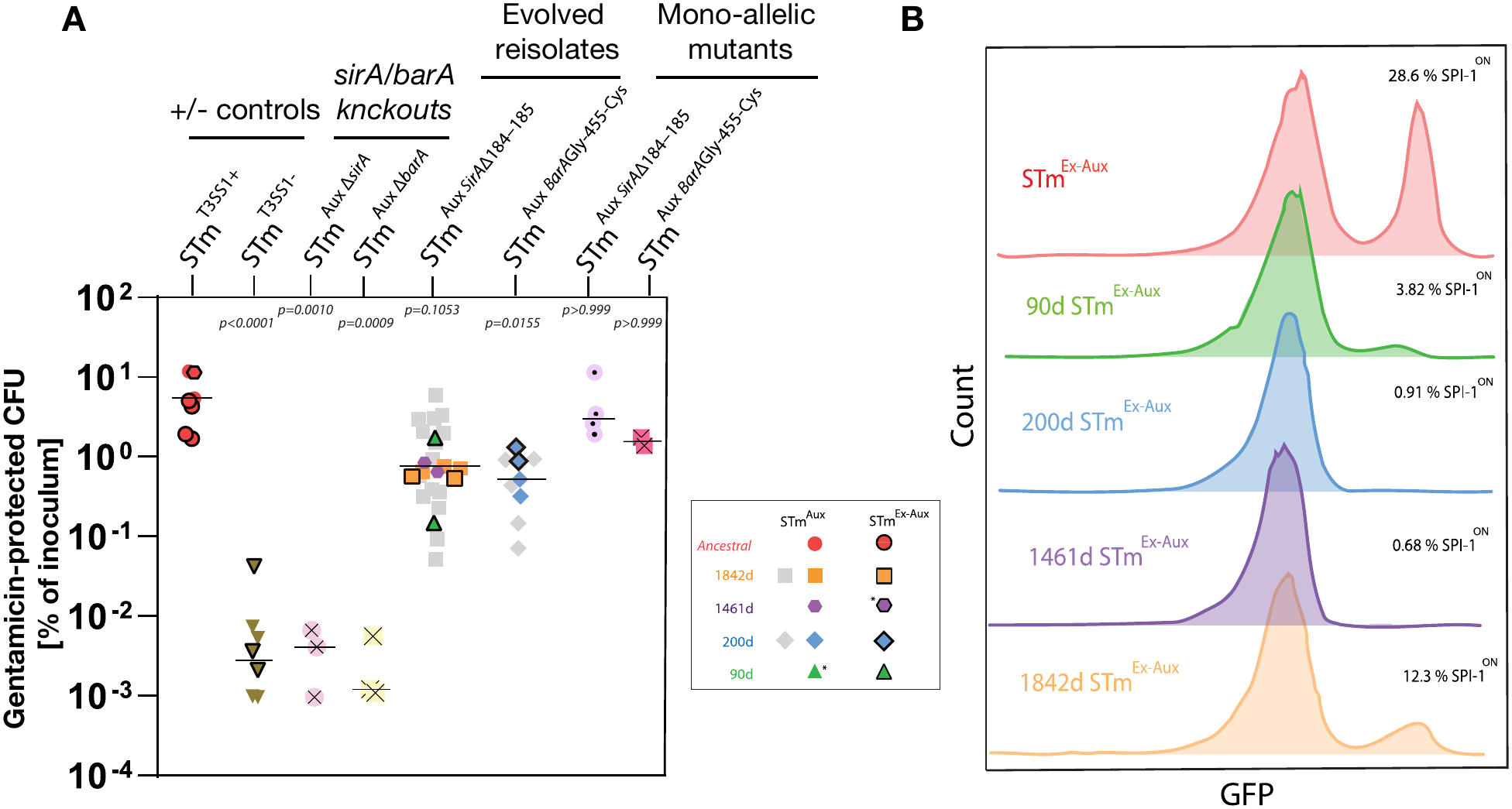
Evolved *barA* and *sirA* mutations modify SPI-1 bistability while preserving epithelial cell invasiveness. **(A)** SPI-1-dependent epithelial cell invasion measured by gentamicin protection assays in HeLa cells. Invasion efficiency is expressed as the percentage of gentamicin-protected CFU relative to the inoculum. SPI-1-competent positive controls (STm^T3SS1+^, red symbols) comprised wild-type STm (hexagon), STm^Aux^ (red circles without outlines), and its ex-auxotrophic derivative STm^Ex-Aux^ (red circles with black outlines), whereas SPI-1-deficient negative controls (STm^T3SS1−,^ inverted brown triangles) consisted of the corresponding Δ*invC* derivatives of STm^Aux^ (symbols without outlines) and STm^Ex-Aux^ (with black outlines). Throughout the figure, black outlines denote strains analysed in the ex-auxotrophic (STm^Ex-Aux^) genetic background. Δ*sirA* and Δ*barA* in-frame deletion mutants in the ancestral STm^Aux^ background, evolved auxotrophic and ex-auxotrophic reisolates, and reconstructed mono-allelic SirA^Δ184–185^ and BarA^Gly455Cys^ mutants in the STm^Aux^ or STm^Ex-Aux^ backgrounds are indicated by the symbols defined in the graphical legend. Selected evolved isolates analysed in (B) subsequent experiments are highlighted in green, blue, purple, and orange as defined in the inset. Each symbol represents the mean of three technical replicates from one independent experiment. Statistical significance was assessed using a Kruskal–Wallis test followed by Dunn’s multiple-comparison test relative to the SPI-1-positive control. **(B)** Representative flow cytometry analysis of SPI-1 expression in selected evolved STm^Ex-Aux^ strains carrying the SPI-1 GFP reporter plasmid pM972 following growth under standard SPI-1-inducing conditions. GFP fluorescence distributions illustrate the relative proportions of phenotypically virulent SPI-1^ON^ and phenotypically avirulent SPI-1^OFF^ subpopulations. Percentages indicate the SPI-1^ON^ fraction. One representative of three independent experiments is shown.

As expected, complete deletion of *sirA* or *barA* abolished cell invasion, phenocopying SPI-1 T3SS-deficient control strains. In contrast, reconstructed strains carrying the evolved *sirA*^c.548-553del^ or the *barA*^c.1363G>T^ alleles retained substantial invasive capacity, demonstrating that neither allele abolishes SPI-1-dependent invasion (**Fig. 4A**). Similar invasion phenotypes were observed in evolved STm^Aux^ re-isolates recovered after 90, 1461, and 1842 days of evolution in LTE-1 and after 200-days in LTE-2, corroborating that also in the evolved strain background *sirA*^c.548-553del^ and the *barA*^c.1363G>T^ mutations preserve SPI-1 dependent cell invasiveness. Although invasiveness tended to be variably reduced relative to the ancestral strain, the effect remained moderate at most and became more pronounced only in late-evolved strains carrying additional adaptive mutations. Consistently, directly *ex vivo* purified populations of evolved STm^Aux^ bacteria from LTE-1 and LTE-2 also displayed reduced, but clearly preserved, SPI-1-dependent HeLa cell invasion (**Supp. Fig. 2**).

These findings suggest that SPI-1 activity is retained although quantitatively altered in long-term evolving strains, prompting us to examine SPI-1 regulation at the single-cell level. Because SPI-1 expression is bistable, generating distinct SPI-1^ON^ and SPI-1^OFF^ subpopulations [1-3], reduced invasion could reflect shifts in the SPI-1^ON^/SPI-1^OFF^ population structure rather than individually reduced SPI-1 expression. Representative evolved STm^Ex-Aux^ re-isolates and control strains were therefore transformed with a transcriptional SPI-1 reporter plasmid carrying a PsicA::gfp fusion [2], grown under standard SPI-1-inducing conditions, and analyzed by flow cytometry (**Fig. 4B**).

Across all evolved strains tested, the proportion of SPI-1^ON^ cells was markedly reduced in *sirA*^c.548-553del^ and *barA*^c.1363G>T^ mutant strains alike (**Fig. 4B**). In contrast, the mean fluorescence intensity of individual SPI-1^ON^ cells remained largely unchanged, indicating preserved SPI-1 transcriptional output at the single-cell level and consistent with the notion that evolved SPI-1^ON^ cells remain individually SPI-1-and invasion-competent.

Notably, genotypes isolated after 200 and 1461 days exhibited SPI-1^ON^ fractions below 1%, approaching the detection limit of this assay, yet retained measurable invasion capacity (**Fig. 4A, B**). This observation is consistent with previous work showing that extremely small SPI-1^ON^ subpopulations can sustain HeLa cell invasion *in vitro*, with as little as 0.17 % of SPI-1^ON^ cells reported to be sufficient for measurable invasion [28]. Thus, within-host evolution of STm^Aux^ reduced the frequency of SPI-1 transcribing cells while at most moderately reducing cell invasion.

Together, these data demonstrate that adaptive *barA*/*sirA* mutations remodel SPI-1 bistable transcriptional regulation by reducing the fraction of SPI-1-expressing cells while preserving the cell invasive functionality of the SPI-1^ON^ subpopulation. These data support a model in which long-term within-host evolution of STm^Aux^ favors quantitative tuning of SPI-1 expression rather than loss of SPI-1 function.

### Monoallelic reconstruction of SirA^Δ184–185^ recapitulates the SPI-1-mdiated virulence phenotype of evolved strains

The experiments above demonstrated that the adaptive *barA*/*sirA* mutations markedly reduce the fraction of SPI-1^ON^ cells while preserving SPI-1-dependent epithelial cell invasion. We next asked whether these regulatory changes affect SPI-1-associated virulence *in vivo*.

Because the auxotrophic STm^Aux^ strain is metabolically unable to sustain pathogenic infection, it induces neither detectable intestinal inflammation nor systemic dissemination despite retaining SPI-1-dependent epithelial invasiveness [9]. Direct assessment of virulence phenotypes therefore requires restoration of the deleted auxotrophy loci. We consequently restored the deleted auxotrophy loci, generating prototrophic “ex-auxotrophic” derivatives (STm^Ex-Aux^) and analyzed them in the streptomycin-pretreated mouse model of invasive non-typhoidal *Salmonella* enterocolitis [33]. In this model, both SPI-1 and SPI-2 contribute to intestinal inflammation, whereas systemic dissemination depends predominantly on SPI-2-mediated intracellular survival [29]. To increase the sensitivity of the model for detecting SPI-1-associated virulence phenotypes, we introduced an additional Δ*ssaV* mutation abolishing SPI-2 function [29].

Using this approach, we examined two independently evolved LTE-1 re-isolates carrying the SirA^Δ184–185^ allele (re-isolated at day 90 and day 1461), together with an isogenic SirA^Δ184–185^ reconstruction in the ancestral strain background (**Fig. 5A**, STm^Aux^ ^SirAΔ184–185^ ^Δ*ssaV*^). An isogenic Δ*sirA* deletion mutant (STm^Aux^ ^Δ*sirA*^ ^Δ*ssaV*^) was included as a control representing complete loss of SirA-dependent virulence regulation.

**Figure 5.**
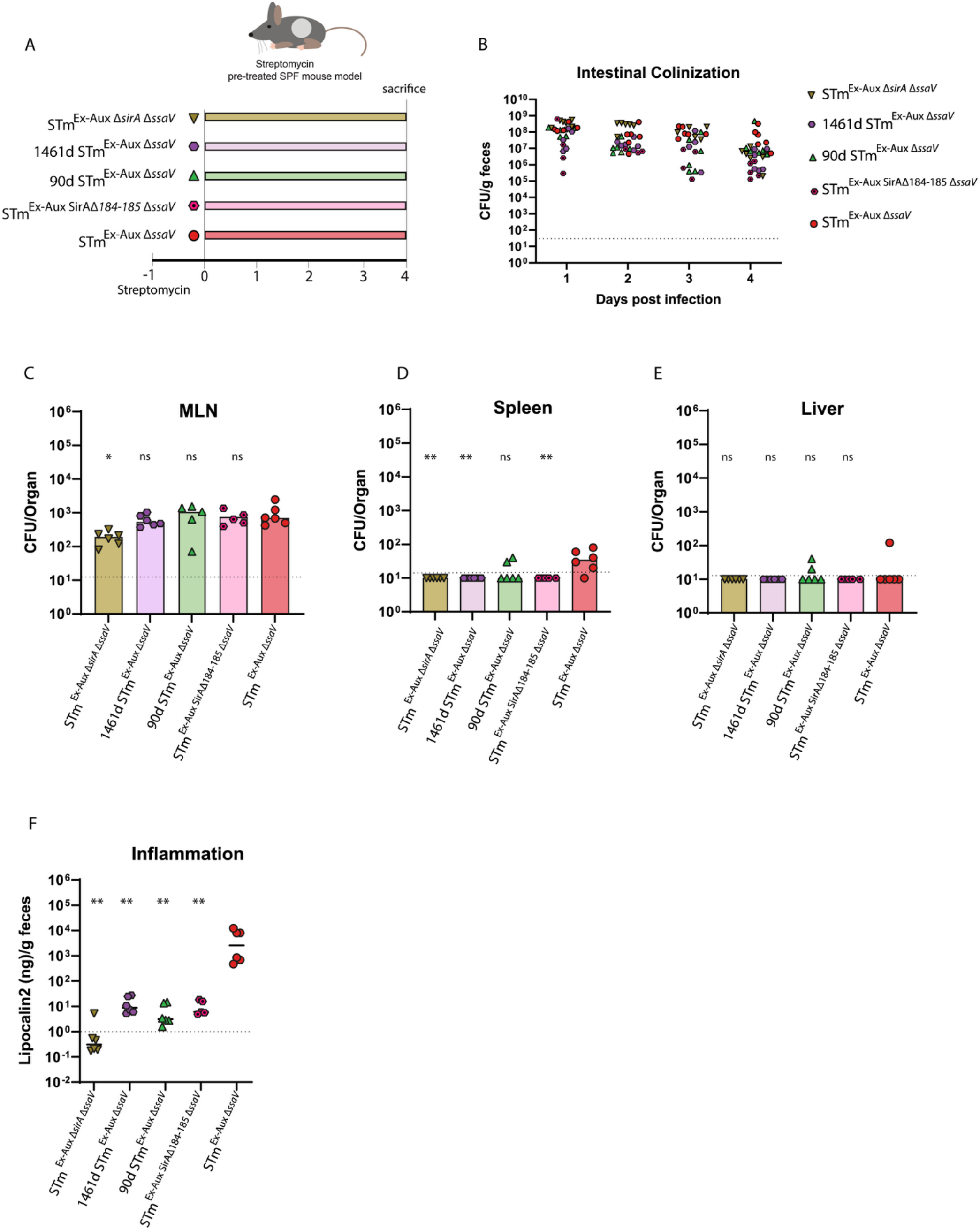
SirA^Δ184-185^ mutation attenuates SPI-1–dependent virulence in a SPI-2–deficient background. **(A)** SPF mice were pretreated with streptomycin (20 mg per mouse) 24 h before oral gavage with 10⁷ CFU of either non-evolved STm^Ex-Aux Δ*ssaV*^ (red circles with black outlines), STm^Ex-Aux^ SirA^Δ184-185 Δ*ssaV*^ (pink hexagon with a central dot), 90d STm^Ex-Aux Δ*ssaV*^ (green triangles with black outlines),1461d STm^Ex-Aux Δ*ssaV*^ (purple hexagons with black outlines) or STm^Ex-Aux Δ*sirA* Δ*ssaV*^ (khaki inverted triangles with black outlines). Fecal samples were collected daily. At 4 days post-infection, mice were sacrificed, and inflammation in the cecum as well as bacterial loads in the mesenteric lymph nodes (MLN), spleens, and livers were quantified by plating on selective media. **(B)** Fecal bacterial populations determined by CFU plating of homogenized fecal samples on selective media. **(C)** Bacterial burden in the mesenteric lymph nodes (MLN) at day 4 post-infection. **(D)** Bacterial burden in the spleens at day 4 post-infection. **(E)** Bacterial burden in the livers at day 4 post-infection. **(F)** Intestinal inflammation at day 4, measured by ELISA quantification of fecal lipocalin-2. Data in panels C-F were analyzed using unpaired, two-tailed Mann–Whitney tests compared with the STm^Ex-Aux^ group. Individual data points are shown, with horizontal bars representing the median. P values: ns, p > 0.05; *p < 0.05; ** p< 0.01; *** p<0.001; **** p<0.0001

All strains efficiently colonized the intestinal lumen throughout the experiment (**Fig. 5B**). Consistent with the SPI-2-deficient genetic background, bacterial burdens in spleen and liver remained at or near the detection limit in all groups (**Fig. 5D, E**). As expected, the Δ*sirA* mutant displayed strongly reduced intestinal virulence-associated phenotypes, characterized by minimal intestinal inflammation and reduced translocation to the mesenteric lymph nodes (MLN) (**Fig. 5C-F**). In contrast, the reconstructed SirA^Δ184–185^ mutant retained measurable virulence activity. Notably, its phenotype closely resembled those of the evolved day-90 and day-1461 re-isolates carrying the same adaptive allele. All three SirA^Δ184–185^ strains induced significantly lower levels of intestinal inflammation than the ancestral STm^Ex-Aux^ ^Δ*ssaV*^ strain while remaining clearly distinguishable from the Δ*sirA* knockout mutant (**Fig. 5F**).

Thus, reconstruction of the SirA^Δ184–185^ allele faithfully reproduced the SPI-1-associated virulence phenotype of evolved SirA^Δ184–185^ strains, demonstrating that this single adaptive mutation is sufficient to confer a partial reduction in SPI-1-associated virulence without abolishing SPI-1 function. Together with the invasion and SPI-1 reporter analyses above (**Fig. 4**), these findings support a model in which adaptive evolution of the BarA/SirA regulatory system reduces the size of the SPI-1-expressing subpopulation while preserving its invasion competence, resulting in quantitatively reduced but clearly preserved virulence-associated host interaction *in vivo*.

### Early evolved *barA/sirA* genotypes retain full virulence in a SPI-2-competent strain background

Although SirA^Δ184–185^ reduced SPI-1-associated virulence in the SPI-2-deficient background (**Fig. 5**), it remained unclear whether this attenuation would remain detectable in a fully virulent genetic background in which both SPI-1 and SPI-2 contribute to disease [29]. We therefore examined representative early evolved STm^Ex-Aux^ re-isolates carrying either SirA^Δ184–185^ (day 90; LTE-1) or BarA^Gly455Cys^ (day 200; LTE-2) in the streptomycin-pretreated mouse model of *Salmonella* enterocolitis [33] without additional SPI-2 attenuation. The day-90 and day-200 isolates were selected because they harbored only a small number of additional mutations beyond SirA^Δ184–185^ and BarA^Gly455Cys^, respectively (**Fig. 6G**). Ancestral STm^Ex-Aux^ served as a virulent positive control, and a SPI-1/SPI-2-deficient STm^Δ*invC*^ ^Δ*ssaV*^ strain served as an avirulent SPI-1 and SPI-2 T3SS-deficient negative control (**Fig. 6A**).

**Figure 6.**
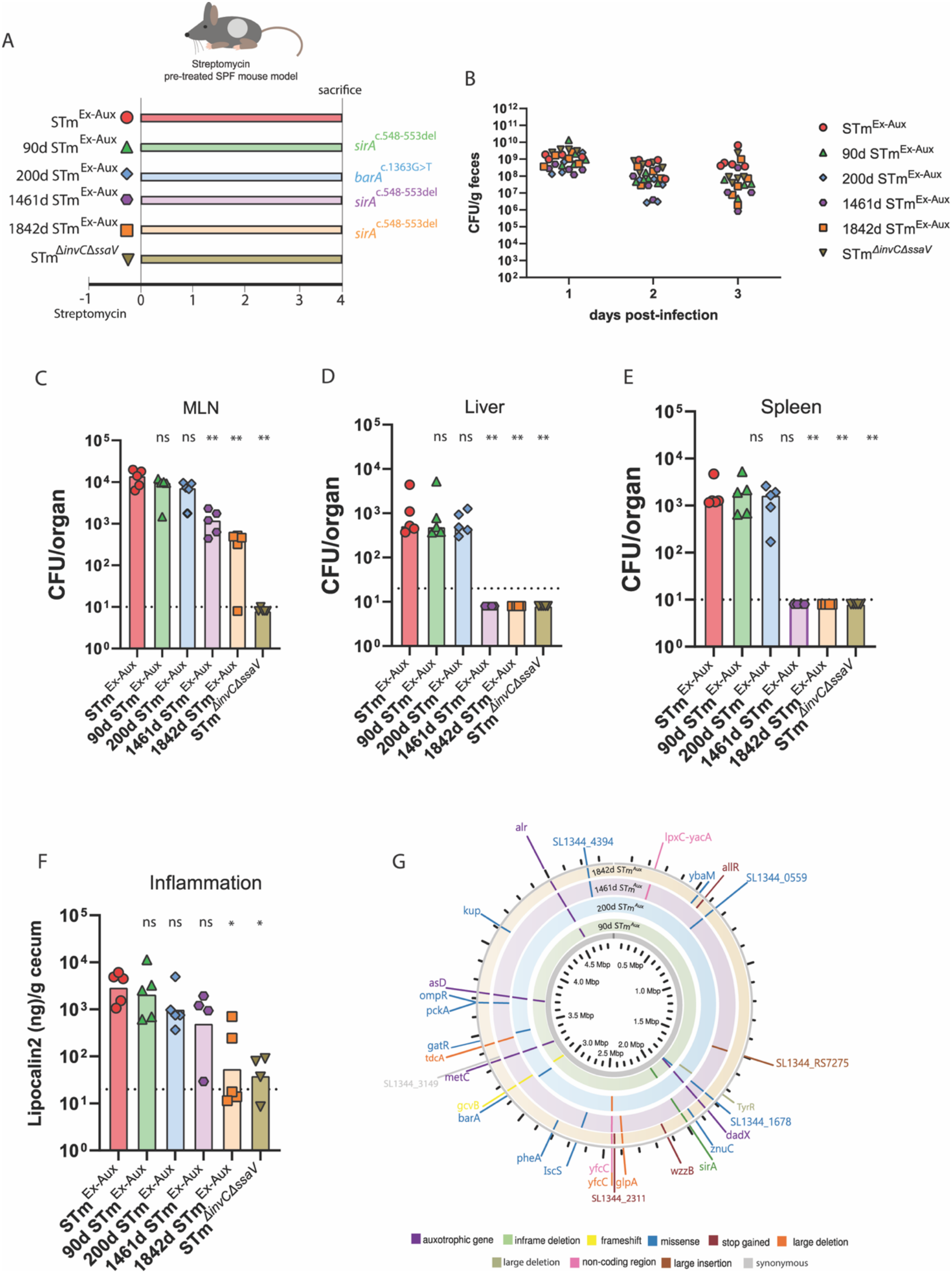
Evolved genotypes harboring SirA^Δ184–185^ and BarA^Gly455Cys^ mutations in a virulence again prototrophic genetic background retain virulence in the iNTS mouse model. **(A)** SPF mice were pretreated with streptomycin (20 mg per mouse) 24 h before oral gavage with 10⁷ CFU of either non-evolved STm^Ex-Aux^ (red circles with black outlines), 90d STm^Ex-Aux^ (green triangles with black outlines), 200d STm^Ex-Aux^ (blue diamonds with black outlines), 1461d STm^Ex-Aux^ (purple hexagons with black outlines), 1842d STm^Ex-Aux^ (orange squares with black outlines) or STm^Δ*invC* Δ*ssaV*^ (khaki inverted triangles with black outlines). Fecal samples were collected daily. At 3 days post-infection, mice were sacrificed, and inflammation in the cecum as well as bacterial loads in the mesenteric lymph nodes (MLN), spleens, and livers were quantified by plating on selective media. Mutations in *sirA* and *barA* are shown for each strain. **(B)** Fecal bacterial populations determined by CFU plating of homogenized fecal samples on selective media. **(C)** Bacterial burden in the mesenteric lymph nodes (MLN) at day 3 post-infection. **(D)** Bacterial burden in the livers at day 3 post-infection. **(E)** Bacterial burden in the spleens at day 3 post-infection. **(F)** Intestinal inflammation at day 3, measured by ELISA quantification of fecal lipocalin-2. **(G)** Circular plot showing genomic mutations identified in 90d STm^Aux^ (green), 200d STm^Aux^ (blue), 1461d STm^Aux^ (purple), and 1842d STm^Aux^ (yellow) following PacBio HiFi sequencing. Mutations are mapped to their corresponding positions in the *Salmonella* genome, and colors indicate the type of mutation. Dotted black line represent the detection limit. Data in panels C–F were analyzed using unpaired, two-tailed Mann–Whitney tests compared with the STm^Ex-Aux^ group. Individual data points are shown, with horizontal bars representing the median. P values: ns, P > 0.05; *P < 0.05; **P < 0.01.

All strains efficiently colonized the intestinal lumen throughout infection (**Fig. 6B**). By day 3 post-infection, both early evolved strains exhibited bacterial burdens in mesenteric lymph nodes (MLN), spleens, and livers comparable to those of the ancestral STm^Ex-Aux^ strain (**Fig. 6C–E**), indicating preserved systemic virulence. Likewise, intestinal inflammation, quantified by fecal lipocalin-2 concentrations, remained comparable to that induced by the ancestral strain (**Fig. 6F**). Thus, despite markedly reduced SPI-1^ON^ populations (**Fig. 4**) and reduced SPI-1-associated virulence in the Δ*ssaV* background (**Fig. 5**), early evolved strains retained full virulence in the conventional SPI-1/SPI-2-competent infection model.

### Long-term evolved STm^Aux^ strain backgrounds exhibit loss of systemic virulence

We extended this analysis to late evolved STm^Ex-Aux^ re-isolates recovered after 1461 and 1842 days of within-host evolution. In contrast to the early evolved strains, both late isolates displayed a profound loss of systemic virulence, characterized by strongly decreased bacterial burdens in spleens and livers (**Fig. 6D, E).** Consistent with this attenuation, bacterial translocation to the mesenteric lymph nodes was also reduced (**Fig. 6C**). However, both strains remained able to induce measurable intestinal inflammation (**Fig. 6F**), indicating that mucosal virulence-associated host interaction was diminished but not abolished.

These findings reveal a progressive divergence of virulence phenotypes during long-term within-host evolution. Whereas early evolved strains carrying adaptive *barA*/*sirA* mutations retained full virulence in the SPI-1/SPI-2-competent infection model, strains recovered after 1461 and 1842 days of evolution exhibited a marked loss of systemic virulence. Notably, this transition occurred without further obvious deterioration of SPI-1-associated phenotypes. The reduction in SPI-1^ON^ population size (**Fig. 4B**), partial preservation of intestinal inflammatory activity (**Fig. 5**), and retained epithelial cell invasion capacity (**Fig. 4A**) remained broadly comparable to those observed in earlier evolved strains. Together, these observations indicate that the adaptive remodeling of SPI-1 bistability remains evolutionarily stable during long-term colonization and is insufficient to explain the subsequent loss of systemic virulence. Instead, additional mutations acquired during prolonged within-host evolution are likely responsible for the emergence of the late hypovirulent phenotype.

Comparative genome analysis identified six mutations shared by the day-1461 and day-1842 isolates but absent from the virulent day-90 strain (**Fig. 6G**). These candidate mutations affect genes involved in membrane-associated regulation and signal transduction (*ompR*) [22,23], central carbon metabolism (*pckA*) [34], glycerol utilization (*glpA-glpQ-glpT* deletion) [24], a large upstream deletion in *tdcA,* a transcriptional activator of the *tdc* operon [35], and two currently poorly characterized proteins encoded by SL1344_0559 and SL1344_4394. Although their individual contributions remain unresolved, these mutations provide plausible candidates for the genetic basis of the late hypovirulent phenotype (**Fig. 6**).

The BarA/SirA regulatory system occupies a central position within the regulatory network governing SPI-1 bistability, acting through the Csr regulatory system to coordinate expression of the HilD virulence regulon in SPI-1^ON^ cells and physiological pathways associated with intestinal luminal colonization in SPI-1^OFF^ cells [34,35]. Because adaptive BarA/SirA variants retained substantial SPI-1 function, and because the BarA/SirA system regulates processes extending beyond SPI-1, we additionally explored whether similar adaptive mutations may also emerge in a HilD-deficient STm^Aux^ strain background.

In parallel with LTE-2, we therefore additionally followed the within-host evolution of an isogenic STm^Aux^ ^Δ*hilD*^ strain during 200 days of colonization of Oligo-MM^12^ mice (**Supp. Fig. 3A**). Whole-genome sequencing of evolved re-isolates revealed partial convergence with the evolutionary trajectories observed in HilD-proficient STm^Aux^ populations (**Supp. Fig. 3C, D**). Most notably, the identical *sirA*^c.548_553del^ allele (SirA^Δ184–185^) arose independently within 200 days of colonization. In addition, an independent *sirA* frameshift mutation likely to cause loss of function (*sirA*^c.175_176insGA^; p.Thr85fs*9) was detected in some isolates. Interestingly, all day-200 STm^Aux^ ^Δ*hilD*^ re-isolates analyzed carried mutations affecting either *sirA* or *ffiZ*, a regulator functionally linked to the BarA/SirA–HilD regulatory network [36-38]. The repeated emergence of mutations affecting components of this regulatory circuit indicates that adaptation repeatedly targets BarA/SirA-associated regulatory pathways even when canonical HilD-mediated SPI-1 regulation is absent.

To exclude the possibility that SirA^Δ184–185^ restores physiologically relevant SPI-1 activity through a HilD-independent pathway, we examined epithelial invasion by evolved and reconstructed STm^Aux^ ^Δ*hilD*^ strains. Neither evolved re-isolates carrying SirA^Δ184–185^ nor an isogenic SirA^Δ184–185^ reconstruction in the ancestral STm^Aux^ ^Δ*hilD*^ background displayed detectable invasion above the background level of the SPI-1-deficient control strain (**Supp. Fig. 3E**). These findings indicate that the adaptive SirA^Δ184–185^ allele does not bypass the requirement for HilD and does not restore SPI-1-dependent epithelial invasion.

Together, these findings demonstrate that recurrent *sirA* evolution occurs even in the absence of HilD and therefore cannot be explained solely by selection acting on canonical HilD-dependent SPI-1 regulation. Because BarA/SirA controls physiological processes extending beyond SPI-1, including pathways involved in luminal fitness and metabolism [39,40], these observations are consistent with selection acting on broader functions of the BarA/SirA regulatory network during long-term intestinal colonization.

## Discussion

How enteric pathogens balance virulence, persistence, and fitness during prolonged host colonization remains a central question in microbial evolution. Virulence programs such as *Salmonella* HilD/SPI-1 are often conceptualized as binary traits that are either maintained because they confer a selective advantage or lost when their fitness costs outweigh their benefits [1,3,4,6,41,42]. Our findings challenge this simplified view. Using a defined model of asymptomatic long-term intestinal colonization, we show that adaptation of STm^Aux^ is dominated not by collapse of virulence functions but by recurrent modification of a global regulatory network controlling the balance between virulence and fitness-associated traits [48,49].

The most striking observation was the repeated and highly parallel evolution of the BarA/SirA regulatory system. Independent long-term evolution experiments repeatedly selected mutations in either *barA* or *sirA*. Remarkably, several of these mutations were identical across independent evolutionary lineages and overlap with variants recently identified in persistent human non-typhoidal *Salmonella* infections [8]. Together with the recurrence of BarA/SirA mutations across independent mouse experiments, human isolates, and previous evolutionary studies [5,7,8], this convergence identifies the BarA/SirA system as a major evolutionary target during prolonged *Salmonella* persistence. Particularly striking was the recurrence of identical or highly similar mutant alleles across independent experimental lineages and persistent human isolates, suggesting that the adaptive landscape available to long-term colonizing *Salmonella* may be constrained. More broadly, these findings are consistent with studies demonstrating that long-term bacterial adaptation frequently proceeds through modification of global regulatory networks rather than structural gene changes alone [32,43,44].

Although BarA/SirA is a central upstream regulator of and required for SPI-1 expression, adaptive evolution did not result in classical SPI-1 loss-of-function phenotypes. Instead, evolved strains retained substantial epithelial invasion capacity and maintained measurable virulence-associated host interaction *in vivo*. Single-cell analyses revealed that adaptive *barA* and *sirA* mutations primarily reduced the proportion of SPI-1^ON^ cells while preserving SPI-1 transcriptional output within the remaining SPI-1-expressing subpopulation. Thus, rather than abolishing SPI-1 function, evolution repeatedly selected for quantitative tuning of SPI-1 bistability.

This mode of adaptation contrasts with evolutionary trajectories described in highly inflammatory settings, where most commonly hilD-deficient SPI-1 “cheater” mutants can transiently expand because they exploit inflammation generated by virulent cooperators [3,5]. In the microbiota-preserving environment studied here, complete loss of SPI-1 function was apparently not favored. Instead, selection repeatedly targeted BarA/SirA, producing genotypes that reduce the population-wide cost of virulence expression while retaining a functional invasion-competent subpopulation. These findings suggest that regulatory fine-tuning may represent an evolutionarily stable solution that balances the benefits and costs of virulence expression during long-term intestinal persistence.

Our results further extend recent observations from persistent human infections. Grote and colleagues reported that persistence-associated BarA/SirA variants exhibit reduced SPI-1 transcription and attenuated virulence phenotypes [8]. The present study confirms and expands these findings by demonstrating that recurrently selected BarA/SirA variants preserve substantial SPI-1 function both *in vitro* and *in vivo*. Importantly, the recurrence of highly similar, and in some cases identical, mutations across experimental evolution and human infection suggest that common selective pressures may shape long-term non-typhoidal *Salmonella* adaptation in both settings [8,50].

A second notable finding emerged after restoration of prototrophy in long-term evolved isolates, which enabled assessment of virulence independently of the auxotrophic attenuation mechanism. Whereas early evolved strains carrying adaptive *barA* or *sirA* alleles retained systemic virulence following reconstruction of the deleted auxotrophy loci, strains evolved for 1461 and 1842 days exhibited strongly reduced systemic dissemination. Notably, this attenuation occurred despite continued evidence of SPI-1-associated host interaction, indicating that the late virulence phenotype cannot be explained by further reduced SPI-1 activity alone. The genetic basis of this attenuation remains unresolved. Candidate mutations – that are absent in the fully virulent day-90 reisolate strain background and shared between day-1461 and day-1842 isolates – affect a known global regulator (*ompR*) [22,23], central metabolism (*pckA*) [34], anaerobic glycerol utilization (*glpA,glpT,glpQ*) [24], transcriptional activator of the tdc operon [35] and several poorly characterized genes, including *SL1344_055S*, which is predicted to encode a σ⁵⁴-associated DNA-binding protein [44].

The independent emergence of the identical SirA^Δ184–185^ allele in a Δ*hilD* background provides additional insight into the selective forces acting on the BarA/SirA regulatory system. Remarkably, adaptive evolution repeatedly targeted the BarA/SirA regulatory system even in the absence of HilD, the downstream master regulator of the SPI-1 regulon. Importantly, the SirA^Δ184–185^ allele did not restore epithelial invasion in the Δ*hilD* background, indicating that it does not bypass the requirement for HilD. Its repeated selection therefore cannot readily be explained by selection acting solely on canonical HilD-dependent SPI-1 expression, particularly because auxotrophic STm^Aux^ is unable to derive detectable pathogenicity from SPI-1-dependent host-interaction [9]. Instead, these observations suggest that adaptive evolution targets broader physiological functions of the BarA/SirA regulatory network, consistent with its established roles in luminal fitness, metabolism, and alternative lifestyle programs [39,40].

From a translational perspective, our findings have implications for the design of live attenuated vaccines [51,53]. Regulatory flexibility, rather than irreversible loss of virulence-associated functions, may contribute to evolutionary stability while preserving desirable immunogenic properties. Notably, adaptive evolution repeatedly targeted regulatory pathways while leaving the engineered auxotrophic attenuation intact throughout more than five years of intestinal colonization. Neither longitudinal phenotypic screening nor genome sequencing of evolved isolates provided evidence of reversion of the engineered auxotrophic phenotype. Preservation of residual SPI-1 function may help maintain mucosal immunogenicity, whereas the continued genetic stability of the auxotrophic lesions supports the robustness of the underlying attenuation strategy. At the same time, these findings emphasize the importance of stable attenuation mechanisms, as host-interaction functions can remain evolutionarily conserved even during extensive adaptation. Our data further suggest that the evolutionary processes associated with persistent non-typhoidal *Salmonella* colonization remains largely intact in STm^Aux^ despite its artificial genetic uncoupling of SPI-1-dependent host interaction from pathogenicity. This observation supports the notion that key evolutionary processes associated with long-term persistence can be preserved in rationally attenuated strains, an encouraging feature for future live-vaccine applications.

Several questions remain unresolved. The molecular mechanisms by which BarA/SirA variants remodel SPI-1 bistability, the physiological traits under selection in the Δ*hilD* background, and the genetic basis of late virulence attenuation all warrant further investigation. In addition, it remains unclear how long-term regulatory adaptation influences the magnitude and quality of mucosal immune responses elicited by persistent vaccine strains.

In summary, our work demonstrates that prolonged within-host evolution of the live vaccine strain STm^Aux^ is not characterized by simple collapse of virulence-associated functions. Instead, adaptation repeatedly targets the BarA/SirA regulatory network, resulting in quantitative remodeling of SPI-1 bistability while preserving substantial functional activity. These findings reveal regulatory adaptation as a major evolutionary route by which persistent Salmonella populations balance fitness and host interaction and provide a framework for understanding long-term pathogen persistence as well as the evolutionary stability of live attenuated bacterial vaccines.

## Material and Methods

### Animal Experiments

All animal experiments were performed in accordance with Swiss Federal law and approved by the Bernese Cantonal Ethical Committee for Animal Experiments (licenses BE85/17, BE100/20, BE28_2021 and BE49_2024). Mice were maintained under strictly controlled conditions at the Clean Mouse Facility (CMF), University of Bern. The ambient temperature was maintained at 23–25°C, with 52–60% relative humidity. Mice were provided ad libitum with autoclaved rodent chow (Kliba-Nafag 3307; Granovit, Switzerland) and autoclaved surgical irrigation water (Baxter Healthcare, ERKF7114).

### Long term colonization and infection experiments

Infection experiments were performed by enteral inoculation of mice with 200 µL of a bacterial suspension in PBS containing either 10⁷ or 10⁸ CFU. Bacterial inocula were prepared as described below.

Gnotobiotic OligoMM^12^ mice had been generated by inoculating germ-free C57BL/6 mice with purified cultures of the murine intestinal bacterial consortium OligoMM^12^. These mice were stably maintained in flexible film isolators under strictly axenic conditions at the Genaxen Clean Mouse Facility (CMF) at the University of Bern.

Two independent long term evolution experiments (LTE-1 and LTE-2) were performed in OligoMM^12^ gnotobiotic mice to monitor *Salmonella* adaptation over time. For LTE-1, mice were kept in open cages under strict bioexclusion inside flexible film isolators for the entire duration of the experiment (up to five years), while for LTE-2, mice were transferred from isolators to IsoCage P bioexclusion cages (Tecniplast, Italy) and maintained there until the end of the experiment. Mice were colonized by inoculation with 10^7^ CFU of STm^Aux^ or STm^Aux^ ^Δ*hilD*^ by gavage, and fecal samples were collected periodically for bacterial population analyses. Breeding pairs were established at specific time points, allowing parallel vertical transfer and adaptation. LTE-1 underwent 11 vertical transfers, whereas LTE-2 included two transfers (**Supp. Fig. 1A–B**).

Specific Pathogen Free (SPF) C57BL/6JRj mice were obtained from Janvier Labs and Charles River and kept in Green-Line IVC cages (Tecniplast, Italy) at the Institute for Infectious Diseases, Bern. For infections using the streptomycin pre-treatment NTS model, SPF mice were pretreated orally with 20 mg of streptomycin 24 hours prior to oral bacterial infection as described [33].

### Bacteria and culture conditions

The bacterial strains and plasmids used in this study are listed in **Supp. Table 1**. All oligonucleotide sequences are listed in **Table 1**.

**Table 1.**
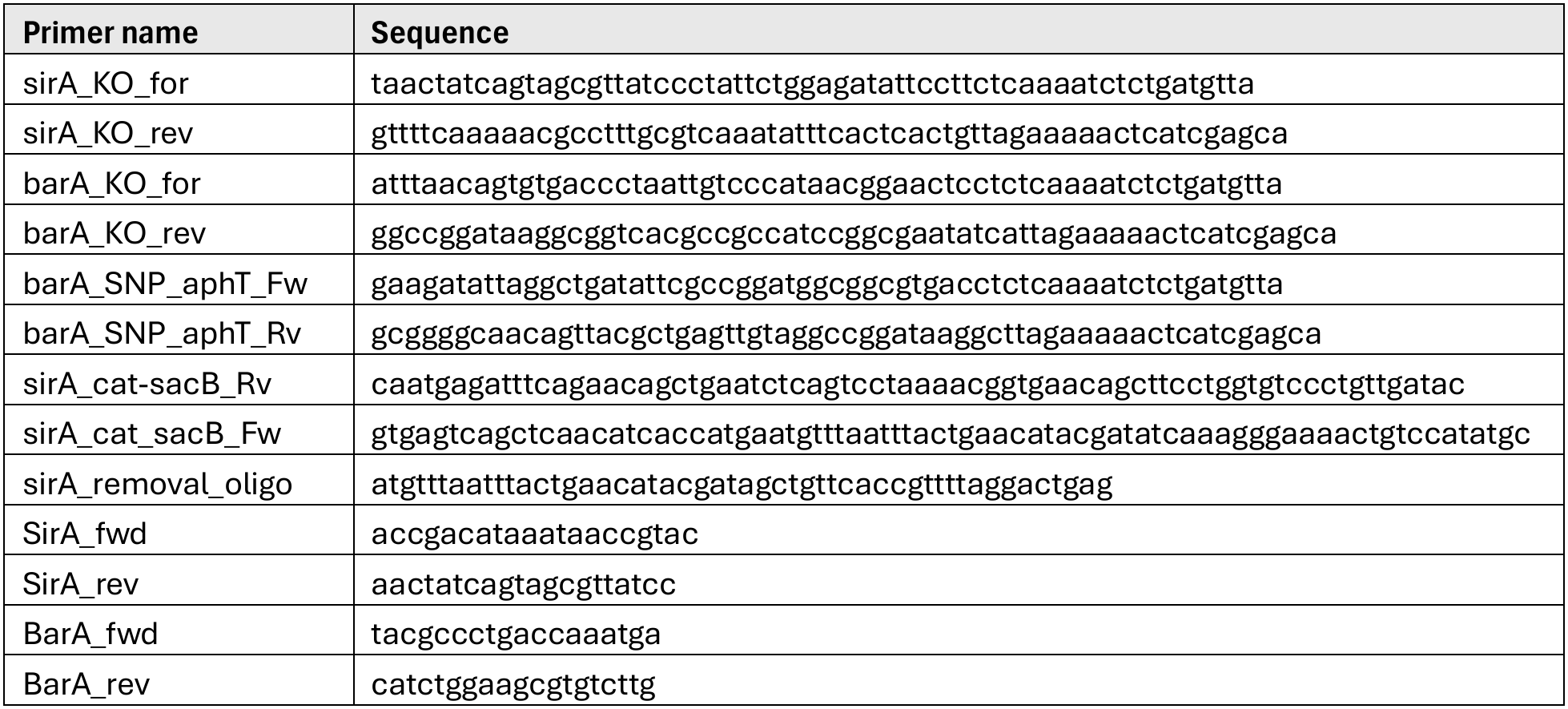
Oligonucleotides used in this study.

Bacteria were routinely cultured in 10 mL Luria-Bertani (LB) medium (Sigma-Aldrich) and incubated at 37°C with shaking at 150 rpm. When required, media were supplemented with ampicillin (100 μg/mL), tetracycline (12.5 μg/mL), kanamycin (50 μg/mL), chloramphenicol (6 μg/mL), streptomycin (50 μg/mL), *meso*-diaminopimelic acid (*m*-DAP; 50 μg/mL), and/or D-alanine (100 μg/mL).

For infection experiments and gentamicin invasion assays, bacterial strains were cultured under SPI-1-inducing conditions in Luria–Bertani (LB) medium supplemented with 0.3 M NaCl, and, when required for auxotrophic strains, D-alanine (100 μg/mL) and meso-diaminopimelic acid (50 μg/mL). Overnight cultures were incubated at 37°C with shaking at 150 rpm for 16 h. For SPI-1 induction, overnight cultures were diluted 1:20 into fresh LB medium supplemented with 0.3 M NaCl and incubated for an additional 4 h under the same conditions before infection. Bacterial cells were then collected by centrifugation (4,816 × g, 15 min, 4°C), washed twice with cold PBS, and resuspended to the desired inoculum concentration.

### Bacterial strains used, generated and isolated in this study

All strains used and generated in this study are listed in **supplementary tables 1 and 2**. The auxotrophic strain HA630 cl.3 (STm^Aux^; SL1344 background; Δ*asd*::tetRA Δ*dadX* Δ*alr* Δ*metC* was constructed from the wild-type strain SB300 by successive rounds of λ-Red recombineering, as previously described [11]. All auxotrophic strains are isogenic to STm^Aux^.

Most genetically engineered bacterial strains used in this study were generated by P22-mediated transduction, as previously described [54]. Isogenic deletion mutants Δ*invC*::*aphT*, Δ*ssaV*::*aphT*, and Δ*hilD*::cat were obtained by P22-mediated transduction using donor strains M736, M738 or HA706, and M315, respectively, according to protocols previously described [1][11]. Similarly, isogenic ex-auxotrophic STm strains (STm^Ex-Aux^) were generated by reintroducing the wild-type alleles of *asd* and either *alr* or *dadX* (see **Supp. Table 1**) into the auxotrophic parental strain background through phage P22 transduction using wild-type STm (strain SB300) as donor strain. Gene deletions and subsequent reinsertions were verified by PCR followed by agarose gel electrophoresis using primer pairs targeting the gene of interest (**Table 1**).

In-frame deletions of *sirA* and *barA* were generated in strains HA2004 and HA2006 by λ-Red-mediated allelic replacement. The *aphT* antibiotic resistance cassette was PCR-amplified using 60-bp primers containing 40-bp homology arms flanking the target gene and 20-bp sequences complementary to the cassette template. For example, sirA deletion was generated using the primers sirA_KO_For and sirA_KO_Rv (see **Table 1**). Purified PCR products (MinElute PCR Purification Kit, Qiagen) were electroporated into strains carrying plasmid pSIM6, which encodes the λ-Red recombination functions, as previously described [55]. Homologous recombination replaced the target coding sequence with the *aphT* cassette while preserving the native stop codon. The resulting mutant alleles were subsequently transferred into the appropriate strain backgrounds by generalized phage transduction, and transductants were selected on LB agar supplemented with kanamycin [56]. Primers used for mutant construction and PCR-based confirmation of genetic modifications are listed in **Table 1**.

STm^Aux^ *barA*^c.1363G>T^ was constructed by inserting a neutral *aphT* cassette into a neutral locus within the P22 transduction range of the gene of interest [56] as previously described [11,57]. The *sirA*^c.548-553del^ mutant was engineered using a two-step allelic exchange strategy based on a *cat–sacB* counter-selectable cassette. The cat–sacB cassette was amplified by PCR from strain CC4231 (provided by Donald Court [57]) using primers containing 50-bp homology arms corresponding to the regions flanking the desired *sirA* mutation and 20-bp sequences homologous to the cassette (sirA_cat-sacB_fwd/Rv). Following λ-Red-mediated integration, recombinant strains were selected on chloramphenicol LB-agar plates and subsequently screened by PCR-based genotyping, with agarose gel electrophoresis used to confirm the expected size of the recombinant allele. The *cat–sacB* cassette was subsequently excised through a second λ-Red recombination event using an oligonucleotide corresponding to the day-1842 STm^Aux^ *sirA* allele. Final mutants were recovered by sucrose counterselection as previously described [57].

The GFP reporter plasmid pM972, encoding GFP under the sicA promoter (sicA::gfp) [2], was introduced into bacterial strains by electroporation. Transformants were selected on LB agar plates supplemented with appropriate antibiotics to maintain plasmid selection.

### Quantification of bacterial loads in organs and feces

Organs, fecal samples, and cecum contents were aseptically collected. Organs were homogenized in 0.5 mL of 0.5% tergitol in PBS, and feces/cecum contents were homogenized in 0.5 mL PBS using a TissueLyzer (TissueLyzer LT, Qiagen; 50 Hz, 3–5 min, with a stainless-steel bead). Resuspended and homogenized liquids were spread in desired dilutions on supplemented MacConkey agar plates with or without D-Ala (100 µg/mL) and *m-*DAP (50 µg/mL), incubated for 16h at 37°C, and followed by counting the colony forming units (CFU). The counts were represented as CFU/organ and plotted using GraphPad Prism version 10.2.1

### Enzyme-linked immunosorbent assays (ELISA)

Total lipocalin-2 concentrations of cecum content and fecal pellets were determined by sandwich ELISA using a commercial mouse lipocalin-2/NGAL ELISA DuoSet (RCD, DY1857), according to the manufacturer’s instructions. Absorbance was measured in a 96-microplate reader (THERMOmax microplate reader) at 405 nm. Lipocalin-2 concentrations were analyzed in Prism 10.2.1 for Windows (GraphPad software Inc) by non-linear regression analysis of the standard curve and expressed as ng lipocalin-2 per gram of sample.

### Cellular Invasion Assays

HeLa (Kyoto) cells were maintained in Dulbecco’s Modified Eagle Medium (DMEM) supplemented with 10% fetal bovine serum (FBS) and incubated at 37 °C with 5% CO₂. For experiments, cells were seeded into 24-well plates and grown for 1 day to reach approx. 80% confluence. Immediately prior to *Salmonella* infection, adherent cells were washed and incubated in Hanks’ Balanced Salt Solution (HBSS). STm strains were cultivated under SPI-1 inducing conditions. 125 mL Erlenmeyer flasks containing 10 mL of LB containing 0.3 M NaCl supplemented with D-Ala (100 µg/mL), *m-*DAP (50 µg/mL) were inoculated from bacterial single colonies and incubated at 37 °C, shaking at 150 rpm for 16 h. Subcultures were prepared by diluting 1:20 into 40 mL of fresh medium and continued incubation under the same conditions until reaching an OD₆₀₀ = 0.35–0.45. Bacteria were then washed in cold PBS. The HeLa cells were infected under HBSS at a multiplicity of infection (MOI) of 1:5 for 50 min, and subsequently extracellular bacteria were eliminated by replacing HBSS with DMEM containing 10% FBS and gentamicin (400 µg/mL) for 30 min. To quantify intracellular (gentamicin-protected) *Salmonella*, HeLa cells were washed once with PBS and lysed in PBS containing 0.1% sodium deoxycholate. The number of gentamicin-protected bacteria was determined by plating the lysate on LB agar supplemented with D-Ala and *m-*DAP.

### STm population-wide genome sequencing analysis

Fecal STm bacteria were enriched by selective plating on MacConkey agar containing streptomycin and appropriate supplements. Following incubation agar plates (containing between 500 and 1000 CFU) STm colonies (spot checked by MALDI-TOF typing to verify identity) were pooled with phosphate-buffered saline (PBS), and the resulting suspension was collected and stores at 4°C until further processing. The plate wash was divided into two aliquots: one aliquot was used for the gentamicin invasion assay (**Supp. Fig. 2**), and the second aliquot was used for genomic DNA extraction (**Fig. 1**). DNA was extracted using either the PureLink™ Microbiome DNA Purification Kit (Invitrogen, USA) or the Wizard® High Molecular Weight (HMW) DNA Extraction Kit (Promega, USA), according to the manufacturers’ instructions.

Genomic DNA quality was assessed by spectrophotometric measurement of A260/280 and A260/230 ratios using a NanoDrop spectrophotometer (Thermo Fisher Scientific, USA). DNA concentrations were determined using the Qubit dsDNA High Sensitivity Assay Kit (Life Technologies, USA) on a Qubit 4.0 fluorometer.

A total of 57 Illumina paired-end DNA libraries (2 × 150 bp) were prepared using the NGS DNA Library Prep Set (Novogene, UK) and sequenced on an Illumina NovaSeq 6000 platform using an S4 flow cell, achieving an average sequencing depth of approximately 100×. Briefly, genomic DNA was randomly fragmented, end-repaired, A-tailed, and ligated to Illumina sequencing adapters. Adapter-ligated fragments were PCR amplified, size-selected, and purified. Library quality and concentration were assessed using the Qubit dsDNA HS assay, quantitative real-time PCR, and a Bioanalyzer for fragment size distribution. Libraries were pooled based on effective concentrations and sequenced at the Novogene UK sequencing facility (Cambridge, UK).

Raw sequencing reads were subjected to quality control using FastQC, followed by adapter removal and quality trimming using Cutadapt. Reads with a Phred quality score below 30 were discarded, and retained reads were trimmed by up to 10 bp based on per-base quality metrics. High-quality reads were aligned to the *Salmonella* enterica subsp. enterica serovar Typhimurium reference genome (GenBank accession NC_016810.1) using Snippy, which incorporates FreeBayes for variant calling. Identified variants were annotated for synonymous and non-synonymous substitutions using SnpEff.

### PacBio high-fidelity genome sequencing analysis

Frozen fecal and cecal samples collected from both long-term experiment mice were plated on diluted MacConkey agar supplemented as described above and containing streptomycin. From each plate, four individual colonies were selected and re-streaked onto fresh MacConkey agar containing the appropriate supplementation and antibiotics. The following day, overnight cultures were prepared as described in the Bacterial culture section. For each isolate, half of the overnight culture was used for PacBio sequencing, while the remaining culture was archived for future analyses.

Overnight cultures of each bacterial strain were grown as described in the Bacterial culture section. Cultures were harvested by centrifugation, and pellets containing no more than 1×10⁹ bacterial cells per sample were used for genomic DNA extraction. High–molecular weight genomic DNA was extracted using the MagAttract® HMW DNA Handbook (Qiagen) according to the manufacturer’s instructions. DNA quantity and integrity were assessed prior to downstream processing to ensure suitability for long-read sequencing.

Purified genomic DNA was mechanically sheared to a target fragment size range of 6–12 kb using a Megaruptor 3 system (Diagenode). DNA dilution for shearing was prepared following the manufacturer’s recommendations to achieve controlled fragmentation. Fragment size distribution and DNA integrity following shearing were assessed using a FEMTO Pulse system (Agilent Technologies), and only samples passing quality control were used for library preparation.

PacBio HiFi libraries were prepared using PacBio’s microbial multiplexing workflow, following the manufacturer’s protocols. Individual samples were barcoded, subjected to library quality control, and pooled in equimolar ratios. A total of 16 samples were multiplexed and sequenced on a single SMRT Cell 25M using the PacBio Revio platform. Circular consensus sequencing (CCS) was performed to generate high-fidelity (HiFi) long-read sequences. Library preparation, quality control, pooling, and sequencing were carried out at the Next Generation Sequencing (NGS) Facility, University of Bern.

Raw sequencing data were processed using PacBio’s standard CCS pipeline to generate HiFi reads. Downstream bioinformatic analyses were performed by Thomas Roder from the Interfaculty Bioinformatics Unit, University of Bern. HiFi reads were mapped to the *Salmonella enterica* serovar Typhimurium SL1344 reference genome (GCF_000210855.2_ASM21085v2_genomic.fna). Single-nucleotide variants and small insertions and deletions were identified using Clair3. Structural variants were identified using Sniffles2.

### Single-cell based SPI-1 expression measurement

The strains STm^Ex-Aux^, 90d STm^Ex-Aux^, 200d STm^Ex-Aux^, 1461d STm^Ex-Aux^, and 1842d STm^Ex-Aux^ were transformed with SPI-1 reporter GFP plasmid pM972 (*sicA::gfp* [2,59]) and grown overnight in in LB containing 0.3 M NaCl, appropriate auxotrophic supplements, and ampicillin (100 µg/mL) under the same SPI-1 inducing conditions as described for the cellular invasion assay. Subcultures diluted 1:20 into fresh medium grown to an OD₆₀₀≈ 0.1.

For flow cytometry analysis, 150 µL of bacterial culture was transferred to a 96-well plate and harvested by centrifugation at 4,700 × g for 7 min at 4 °C. The supernatant was discarded, and pellets were washed once with sterile-filtered PBS, centrifuged, and resuspended in PBS containing Pyronin Y (0.5 µg/mL). After a second centrifugation step, bacterial pellets were fixed in 2% formaldehyde (FA) in PBS (ROTI®Histofix, Carl Roth, Art.-Nr. P087.4) and recorded using a Cytoflex S flow cytometer. Flow cytometry data were processed with FlowJo software (v10.6.1), applying gating strategies to exclude debris and doublets.

### Statistics

Statistical analysis was performed as indicated in Figure legends using Graph Pad Prism 10.2.1.

## Data availability

The raw sequencing data generated in this study are being prepared for deposition in a public sequence repository. Repository accession numbers will be provided in an updated version of this preprint. The sequencing data will be made publicly available no later than publication of the peer-reviewed article.

## Author contributions

L.L.C. performed most experiments, curated and analyzed the data, generated the figures, and contributed to manuscript writing. D.T., O.S., and L.B. planned and conducted the long-term colonization studies. L.L.C., O.S., and M.M conducted all remaining animal studies, virulence studies, mutant strain construction, and invasion assays. D.T. carried out shotgun Illumina sequencing and bioinformatic variant-calling analyses. M.M. and L.L.C. performed PacBio HiFi genome sequencing analyses and downstream bioinformatic characterization of strain re-isolates. D.T and F.A. carried out population-level invasion assays. F.O.-B. carried out flow cytometry analyses. K.F. assisted wet-lab and animal experiments. L.L.C., O.S., and D.T. contributed to experimental design. L.L.C., L.B., and O.S. genetically engineered bacterial strains. L.L.C., O.S., and S.H. curated, integrated, analyzed, and interpreted the experimental and genomic datasets. L.L.C. and S.H. performed the comparative evolutionary analyses and developed the conceptual framework of the study. S.H. conceived, supervised and acquired funding for the study. S.H. and L.L.C. wrote the manuscript. All authors reviewed and edited the manuscript and approved the final version.

## Acknowledgments

We thank Wolf-Dietrich Hardt for providing bacterial strains, genetic tools, and valuable scientific input. We are grateful to Emma Slack for helpful discussions and support throughout the project. We thank Donald Court for providing recombineering tools and expertise. We acknowledge the staff of the Clean Mouse Facility for excellent animal husbandry and technical assistance. We also thank the Genaxen Foundation for their support. Sequencing was performed at the NGS Core Facility, and we thank the Bioinformatics Center for assistance with data processing and analysis. This work was funded by grants from the European Research Council (erc.europa.eu) under the European Union’s Seventh Framework Programme (FP/2007-2013), ERC Grant Agreement n. 281904 (ERC-2013-StG-281904) and the Swiss National Science Foundation (grant 31003A_169791/1; NCCR Microbiomes grants 51NF40_180575 and 51NF40_ 225148). D.T. and S.H. received additional support from the Helmut-Horten Stiftung (helmut-horten-stiftung.org).

