## Supplementary Figures 1-3 and Table 1 for "Within-host evolution of an avirulent *Salmonella* vaccine strain reveals regulatory tuning of virulence functionality without loss of function"

### Supplementary Material

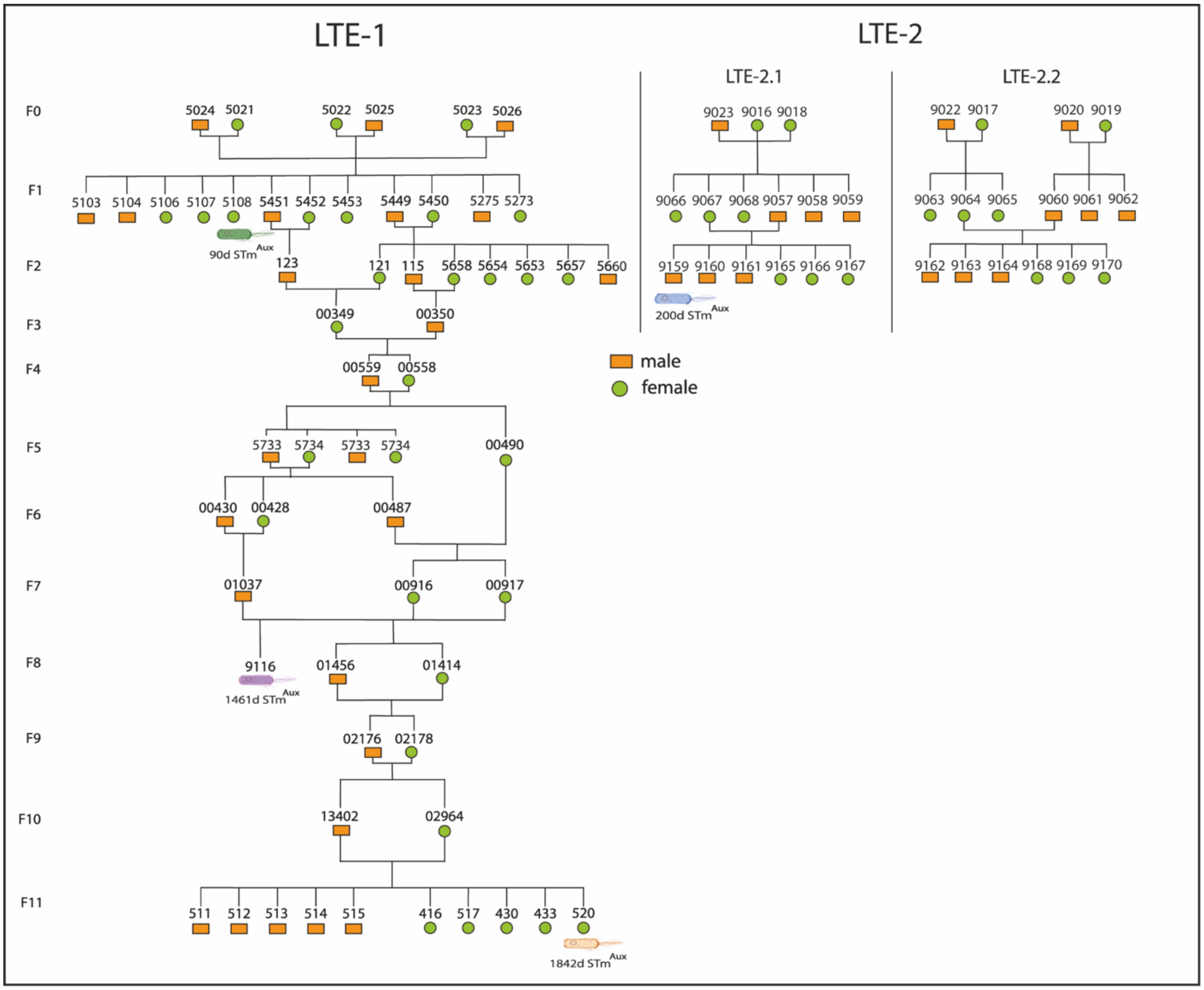


**Supplementary Figure 1. Longitudinal evolution of STm$^{Aux}$ across generations in the Oligo-MM$^{12}$ mouse model.** Within-host evolutionary lineages of LTE-1 and LTE-2, including the independent LTE-2.1 and LTE-2.2 lineages, during intestinal colonization in Oligo-MM$^{12}$ mice. Selected evolved strains analysed for *in vivo* virulence are indicated below the corresponding mouse ID. Number of vertical transfers is shown on the left (F0 to F11).

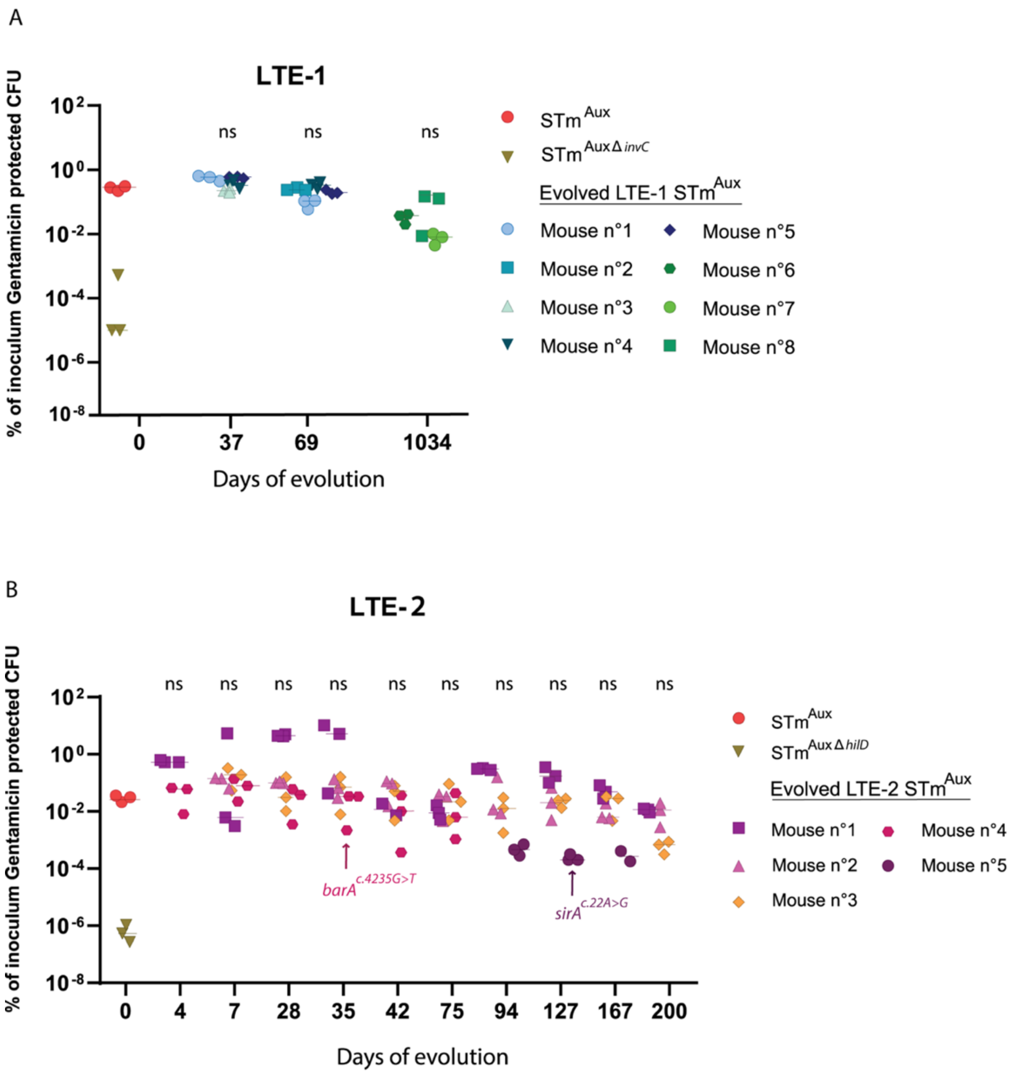


**Supplementary Figure 2. Convergent evolution of BarA/SirA in multiple STm$^{Aux}$ lineages does not abolish SPI-1-dependent invasiveness. (A) and (B)** SPI-1 T3SS–dependent invasion of epithelial HeLa cells by evolved STm$^{Aux}$ populations recovered from plate washes at different time points during LTE-1 **(A)** and LTE-2 **(B)**, quantified by gentamicin protection assay in HeLa cells. Invasion efficiency is expressed as the percentage of gentamicin-protected intracellular CFU relative to the initial inoculum. Each mouse was tested once with three technical replicates; bars indicate the mean of these triplicates. The ancestral STm$^{Aux}$ strain was used as a positive control (red circles), while STm$^{Aux\ \Delta invC}$ and STm$^{Aux\ \Delta hilD}$ served as negative controls (khaki inverted triangles) for LTE-1 (**A**) and LTE-2 (**B**), respectively. Statistical significance was assessed using a Kruskal–Wallis test followed by Dunn's multiple comparisons test, relative to STm$^{Aux}$ ($p$< 0.05).

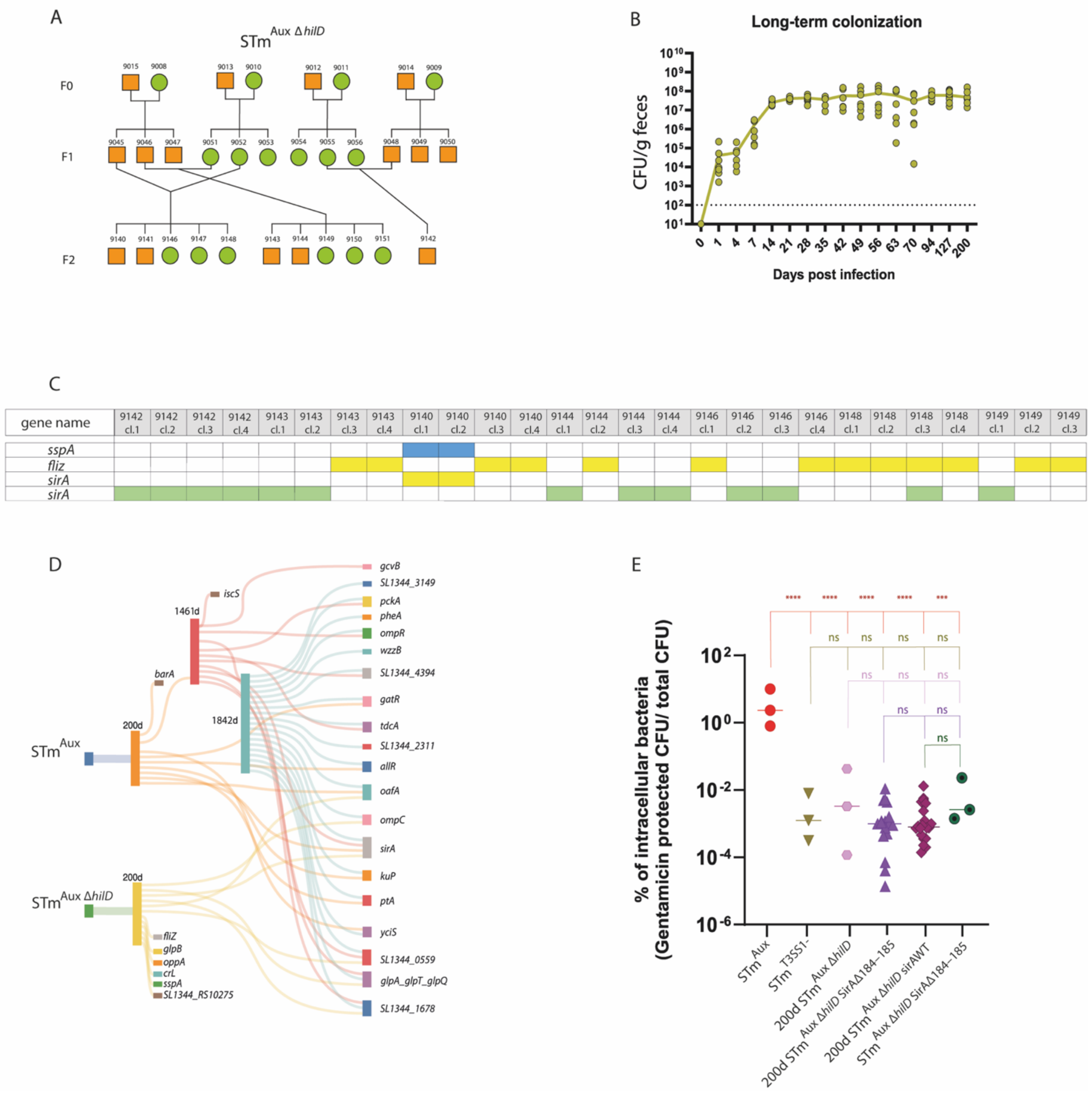


**Supplementary Figure 3. SirA$^{\Delta184–185}$ emerges in evolved STm$^{\text{Aux } \Delta hilD}$ without restoring SPI-1-dependent invasiveness (A)** Within-host phylogeny of STm$^{\text{Aux } \Delta hilD}$ populations recovered from Oligo-MM[12] mice across two successive host passages. (**B)** Long-term intestinal colonization dynamics of STm$^{\text{Aux } \Delta hilD}$ in Oligo-MM[12] mice, quantified over the course of the experiment. (**C)** Distribution of virulence-associated genes [58] in 200-day evolved STm$^{\text{Aux } \Delta hilD}$ re-isolates from seven mice. Between three and four re-isolates per mouse were selected and analysed using PacBio HiFi long-read sequencing. **(D)** Sankey diagram illustrating parallel genomic evolution in STm$^{\text{Aux}}$ and STm$^{\text{Aux } \Delta hilD}$ lineages across three independent long-term evolution experiments, with three sampling evolution time points (200d, 1461d, and 1842d). Mutated genes are represented by colored squares connected to the days of evolution in which they occured. Sankey diagram was generated using sankeymatic.com. **(E)** Gentamicin protection assays comparing HeLa cell invasion by STm$^{\text{Aux}}$ (red circles, positive control), negative control STm$^{\text{Aux } \Delta invC}$ (STm$^{\text{Aux } \Delta invC}$*,* inverted triangles), and various 200-day evolved STm$^{\text{Aux } \Delta hilD}$ re-isolates. Re-isolates are grouped according to their *sirA* genotype being SirA$^{\Delta184–18}$ (200d STm$^{\text{Aux } \Delta hilD \text{ SirA}\Delta184–185}$) or wild-type (200d STm$^{\text{Aux } \Delta hilD \; sirA\text{WT}}$). To specifically assess the impact of the SirA$^{\Delta184–185}$ mutation on invasion, the single mutation was reconstructed in the ancestral STm$^{Aux\ \Delta hilD}$ strain background (green circles). Each data point represents the mean of three technical replicates from one independent experiment. Statistical analysis in panel **E** was performed using one-way ANOVA with Šídák's multiple-comparisons test relative to STm$^{Aux}$ (red), STm$^{Aux\ \Delta invC}$ (khaki), 200 wp STm$^{Aux\ \Delta hilD}$ (pink), 200d STm$^{Aux\ \Delta hilD\ SirA\Delta 184–185}$ (purple) or 200d STm$^{Aux\ \Delta hilD\ sirAWT}$ (magenta). P values: ns, P > 0.05; *P < 0.05; **P < 0.01; ***P < 0.001; ****P < 0.0001.

#### Supplementary Table 1. Bacterial strains, isolates and plasmids.

| Acronym | Strain name | Strain Background /parental strain | Relevant genotype | Resistances/ Auxotrophies | Figure in this study | Origin or Reference |
|---|---|---|---|---|---|---|
| M736 | M736 | ATCC 14028s derivate IR715 | Δ*invC*::aphT | Nal, Kan$^R$ | / | [60] |
| M738 | M738 | ATCC 14028s derivate IR715 | Δ*ssaV*::cat | Nal, Cam$^R$ | / | [60] |
| HA706 | HA706 | SB300 | Δ*ssaV*::cat | Cam$^R$ | / | [9] |
| M315 | M315 | SB300 | Δ*hilD*::cat | Cam$^R$ | / | [1] |
| pM972 | | | | Amp$^R$ | Fig. 4B | [59] |
| STm | SB300 | SB300 | Wild-type, STm SL1344 | Str$^R$ | Fig. 4A | [57] |
| STm$^{Aux}$ | HA630 cl.3 | SB300 | Δ*asd*::tetRA Δ*dadX* Δ*alr* Δ*metC* | Auxotroph Tet$^R$, Str$^R$ | Fig. 1<br>Fig. 4A<br>Supp. Fig. 1<br>Supp. Fig. 2<br>Supp. Fig. 3D, E | [9] |
| STm$^{Aux}$ | HA630 cl.3 1K | HA630 cl.3 | Δ*asd*::tetRA Δ*dadX* Δ*alr* Δ*metC* | Auxotroph Tet$^R$, Str$^R$, Kan$^R$ | Fig. 2 | [9] |
| STm$^{Aux}$ | HA630 cl.3 1C | HA630 cl.3 | Δ*asd*::tetRA Δ*dadX* Δ*alr* Δ*metC* | Auxotroph Tet$^R$, Str$^R$, Cam$^R$ | Fig. 2 | [9] |
| STm$^{Ex\text{-}Aux}$ | HA630 $^{Ex\text{-}Aux}$ | HA630 cl.3 | Δ*dadX* Δ*metC* | Ex-Auxotroph Str$^R$ | Fig. 4A, B<br>Fig. 6 | this study |
| STm$^{Ex\text{-}Aux\ \Delta ssaV}$ | STm$^{Ex\text{-}Aux\ \Delta ssaV}$ | STm$^{Ex\text{-}Aux}$ | Δ*dadX* Δ*metC* Δ*ssaV*::cat | Ex-Auxotroph Str$^R$, Cam$^R$ | Fig. 5 | this study |
| STm$^{Aux\ \Delta invC}$ | HA700 | HA630 cl.3 | Δ*asd*::tetRA Δ*dadX* Δ*alr* Δ*metC* Δ*invC*::aphT | Auxotroph Tet$^R$, Str$^R$, Kan$^R$ | Fig. 4A<br>Supp. Fig. 2B<br>Supp. Fig. 3E | [9] |

| | | | | | | |
|---|---|---|---|---|---|---|
| STm$^{\text{Ex-Aux Δ}invC}$ | HA2045 | HA700 | Δ*dadX* Δ*metC* Δ*invC*::aphT | Ex-Auxotroph Str$^R$, Kan$^R$ | Fig. 4A | this study |
| STm$^{\text{Aux Δ}hilD}$ | HA902 | HA630 cl.3 | Δ*asd*::tetRA Δ*dadX* Δ*alr* Δ*metC* Δ*hilD*::cat | Str$^R$, Cam$^R$ | Supp. Fig. 2C Supp. Fig. 3A-C | this study |
| STm$^{\Delta invC\ \Delta ssaV}$ | HA705 | SB300 | Δ*invC*::aphT Δ*ssaV*::cat | Str$^R$, Kan$^R$, Cam$^R$ | Fig. 6 | [9] |
| STm$^{\text{Ex-Aux Δ}sirA\ \Delta ssaV}$ | STm$^{\text{Ex-Aux Δ}sirA\ \Delta ssaV}$ | HA2046 | Δ*dadX* Δ*metC* Δ*sirA*::aphT Δ*ssaV*::cat | Str$^R$, Kan$^R$, Cam$^R$ | Fig. 5 | this study |
| STm$^{\text{Aux SirAΔ184-185}}$ | HA2051 | HA630 cl.3 | Δ*asd*::tetRA Δ*dadX* Δ*alr* Δ*metC* | Auxotroph Tet$^R$, Str$^R$ | Fig. 4A | this study |
| STm$^{\text{Ex-Aux SirAΔ184-185 Δ}ssaV}$ | HA2051$^{\text{Ex-Aux}}$ | HA2051 | Δ*dadX* Δ*metC* Δ*ssaV*::cat | Ex-Auxotroph Str$^R$, Cam$^R$ | Fig. 5 | this study |
| STm$^{\text{ΔsirA}}$ | HA2004 | HA630 cl.3 | Δ*asd*::tetRA Δ*dadX* Δ*alr* Δ*metC* Δ*sirA*::aphT | Auxotroph Tet$^R$, Str$^R$, Kan$^R$ | Fig. 4A | this study |
| STm$^{\text{ΔbarA}}$ | HA2006 | HA630 cl.3 | Δ*asd*::tetRA Δ*dadX* Δ*alr* Δ*metC* Δ*barA*::aphT | Auxotroph Tet$^R$, Str$^R$, Kan$^R$ | Fig. 4A | this study |
| STm$^{\text{Aux BarAGly455Cys}}$ | HA2054 | HA630 cl.3 | Δ*asd*::tetRA Δ*dadX* Δ*alr* Δ*metC* *barA*G455C::aphT | Auxotroph Tet$^R$, Str$^R$, Kan$^R$ | Fig. 4A | this study |
| 90d STm$^{\text{Aux}}$ | 90d STm$^{\text{Aux}}$ | HA630 cl.3 | Δ*asd*::tetRA Δ*dadX* Δ*alr* Δ*metC* | Auxotroph Tet$^R$, Str$^R$ | Fig. 1A | this study |
| 90d STm$^{\text{Ex-Aux}}$ | HA753 | 90d STm$^{\text{Aux}}$ | Δ*dadX* Δ*metC* | Ex-Auxotroph Str$^R$ | Fig. 4A, B Fig. 6 | this study |
| 90d STm$^{\text{Ex-Aux Δ}ssaV}$ | 90d STm$^{\text{Ex-Aux Δ}ssaV}$ | HA753 | Δ*dadX* Δ*metC* Δ*invC*::cat | Ex-Auxotroph Str$^R$, Cam$^R$ | Fig. 5 | this study |
| 200d STm$^{\text{Aux}}$ | 200d 9159 cl.1 | HA630 cl.3 | Δ*asd*::tetRA Δ*dadX* Δ*alr* Δ*metC* | Auxotroph Tet$^R$, Str$^R$ | Fig. 4A- STm$^{\text{Aux BarAGly455Cys}}$ | this study |
| 200d STm$^{\text{Aux}}$ | 200d 9159 cl.2 | HA630 cl.3 | Δ*asd*::tetRA Δ*dadX* Δ*alr* Δ*metC* | Auxotroph Tet$^R$, Str$^R$ | Fig. 4A-STm$^{\text{Aux BarAGly455Cys}}$ | this study |
| 200d STm$^{\text{Aux}}$ | 200d 9159 cl.3 | HA630 cl.3 | Δ*asd*::tetra Δ*dadX* Δ*alr* Δ*metC* | Auxotroph Tet$^R$, Str$^R$ | Fig. 4A-STm$^{\text{Aux BarAGly455Cys}}$ | this study |
| 200d STm$^{\text{Ex-Aux}}$ | 200d 9159 cl.3 $^{\text{Ex-Aux}}$ | HA630 cl.3 | Δ*dadX* Δ*metC* | Ex-Auxotroph Str$^R$ | Fig. 4A, B Fig. 6 | this study |
| 200d STm$^{\text{Aux}}$ | 200d 9159 cl.4 | HA630 cl.3 | Δ*asd*::tetRA Δ*dadX* Δ*alr* Δ*metC* | Auxotroph Tet$^R$, Str$^R$ | Fig. 4A- STm$^{\text{Aux BarAGly455Cys}}$ | this study |
| 200d STm$^{\text{Aux}}$ | 200d 9161 cl.1 | HA630 cl.3 | Δ*asd*::tetRA Δ*dadX* Δ*alr* Δ*metC* | Auxotroph Tet$^R$, Str$^R$ | Fig. 4A- STm$^{\text{Aux BarAGly455Cys}}$ | this study |

| | | | | | | |
|---|---|---|---|---|---|---|
| 200d $STm^{Aux}$ | 200d 9161 cl.2 | HA630 cl.3 | Δ*asd*::tetRA Δ*dadX* Δ*alr* Δ*metC* | Auxotroph $Tet^R$, $Str^R$ | Fig. 4A- $STm^{Aux}$ BarAGly455Cys | this study |
| 200d $STm^{Aux\ \Delta hilD}$ | 200d HA902 9142 cl.1 | HA902 | Δ*asd*::tetRA Δ*dadX* Δ*alr* Δ*metC* Δ*hilD*::cat | Auxotroph $Tet^R$, $Str^R$, $Cam^R$ | Supp. Fig. 3C | this study |
| 200d $STm^{Aux\ \Delta hilD}$ | 200d HA902 9142 cl.2 | HA902 | Δ*asd*::tetRA Δ*dadX* Δ*alr* Δ*metC* Δ*hilD*::cat | Auxotroph $Tet^R$, $Str^R$, $Cam^R$ | Supp. Fig. 3C | this study |
| 200d $STm^{Aux\ \Delta hilD}$ | 200d HA902 9142 cl.3 | HA902 | Δ*asd*::tetRA Δ*dadX* Δ*alr* Δ*metC* Δ*hilD*::cat | Auxotroph $Tet^R$, $Str^R$, $Cam^R$ | Supp. Fig. 3C | this study |
| 200d $STm^{Aux\ \Delta hilD}$ | 200d HA902 9142 cl.4 | HA902 | Δ*asd*::tetRA Δ*dadX* Δ*alr* Δ*metC* Δ*hilD*::cat | Auxotroph $Tet^R$, $Str^R$, $Cam^R$ | Supp. Fig. 3C | this study |
| 200d $STm^{Aux\ \Delta hilD}$ | 200d HA902 9143 cl.1 | HA902 | Δ*asd*::tetRA Δ*dadX* Δ*alr* Δ*metC* Δ*hilD*::cat | Auxotroph $Tet^R$, $Str^R$, $Cam^R$ | Supp. Fig. 3C | this study |
| 200d $STm^{Aux\ \Delta hilD}$ | 200d HA902 9143 cl.2 | HA902 | Δ*asd*::tetRA Δ*dadX* Δ*alr* Δ*metC* Δ*hilD*::cat | Auxotroph $Tet^R$, $Str^R$, $Cam^R$ | Supp. Fig. 3C | this study |
| 200d $STm^{Aux\ \Delta hilD}$ | 200d HA902 9143 cl.3 | HA902 | Δ*asd*::tetRA Δ*dadX* Δ*alr* Δ*metC* Δ*hilD*::cat | Auxotroph $Tet^R$, $Str^R$, $Cam^R$ | Supp. Fig. 3C | this study |
| 200d $STm^{Aux\ \Delta hilD}$ | 200d HA902 9143 cl.4 | HA902 | Δ*asd*::tetRA Δ*dadX* Δ*alr* Δ*metC* Δ*hilD*::cat | Auxotroph $Tet^R$, $Str^R$, $Cam^R$ | Supp. Fig. 3C | this study |
| 200d $STm^{Aux\ \Delta hilD}$ | 200d HA902 9140 cl.1 | HA902 | Δ*asd*::tetRA Δ*dadX* Δ*alr* Δ*metC* Δ*hilD*::cat | Auxotroph $Tet^R$, $Str^R$, $Cam^R$ | Supp. Fig. 3C | this study |
| 200d $STm^{Aux\ \Delta hilD}$ | 200d HA902 9140 cl.2 | HA902 | Δ*asd*::tetRA Δ*dadX* Δ*alr* Δ*metC* Δ*hilD*::cat | Auxotroph $Tet^R$, $Str^R$, $Cam^R$ | Supp. Fig. 3C | this study |
| 200d $STm^{Aux\ \Delta hilD}$ | 200d HA902 9140 cl.3 | HA902 | Δ*asd*::tetRA Δ*dadX* Δ*alr* Δ*metC* Δ*hilD*::cat | Auxotroph $Tet^R$, $Str^R$, $Cam^R$ | Supp. Fig. 3C | this study |
| 200d $STm^{Aux\ \Delta hilD}$ | 200d HA902 9140 cl.4 | HA902 | Δ*asd*::tetRA Δ*dadX* Δ*alr* Δ*metC* Δ*hilD*::cat | Auxotroph $Tet^R$, $Str^R$, $Cam^R$ | Supp. Fig. 3C | this study |
| 200d $STm^{Aux\ \Delta hilD}$ | 200d HA902 9144 cl.1 | HA902 | Δ*asd*::tetRA Δ*dadX* Δ*alr* Δ*metC* Δ*hilD*::cat | Auxotroph $Tet^R$, $Str^R$, $Cam^R$ | Supp. Fig. 3C | this study |
| 200d $STm^{Aux\ \Delta hilD}$ | 200d HA902 9144 cl.2 | HA902 | Δ*asd*::tetRA Δ*dadX* Δ*alr* Δ*metC* Δ*hilD*::cat | Auxotroph $Tet^R$, $Str^R$, $Cam^R$ | Supp. Fig. 3C | this study |
| 200d $STm^{Aux\ \Delta hilD}$ | 200d HA902 9144 cl.3 | HA902 | Δ*asd*::tetRA Δ*dadX* Δ*alr* Δ*metC* Δ*hilD*::cat | Auxotroph $Tet^R$, $Str^R$, $Cam^R$ | Supp. Fig. 3C | this study |
| 200d $STm^{Aux\ \Delta hilD}$ | 200d HA902 9144 cl.4 | HA902 | Δ*asd*::tetRA Δ*dadX* Δ*alr* Δ*metC* Δ*hilD*::cat | Auxotroph $Tet^R$, $Str^R$, $Cam^R$ | Supp. Fig. 3C | this study |
| 200d $STm^{Aux\ \Delta hilD}$ | 200d HA902 9146 cl.1 | HA902 | Δ*asd*::tetRA Δ*dadX* Δ*alr* Δ*metC* Δ*hilD*::cat | Auxotroph $Tet^R$, $Str^R$, $Cam^R$ | Supp. Fig. 3C | this study |

| | | | | | | |
|---|---|---|---|---|---|---|
| 200d STm$^{Aux\ \Delta hilD}$ | 200d HA902 9146 cl.2 | HA902 | Δ*asd*::tetRA Δ*dadX* Δ*alr* Δ*metC* Δ*hilD*::cat | Auxotroph Tet$^R$, Str$^R$, Cam$^R$ | Supp. Fig. 3C | this study |
| 200d STm$^{Aux\ \Delta hilD}$ | 200d HA902 9146 cl.3 | HA902 | Δ*asd*::tetRA Δ*dadX* Δ*alr* Δ*metC* Δ*hilD*::cat | Auxotroph Tet$^R$, Str$^R$, Cam$^R$ | Supp. Fig. 3C | this study |
| 200d STm$^{Aux\ \Delta hilD}$ | 200d HA902 9146 cl.4 | HA902 | Δ*asd*::tetRA Δ*dadX* Δ*alr* Δ*metC* Δ*hilD*::cat | Auxotroph Tet$^R$, Str$^R$, Cam$^R$ | Supp. Fig. 3C | this study |
| 200d STm$^{Aux\ \Delta hilD}$ | 200d HA902 9148 cl.1 | HA902 | Δ*asd*::tetRA Δ*dadX* Δ*alr* Δ*metC* Δ*hilD*::cat | Auxotroph Tet$^R$, Str$^R$, Cam$^R$ | Supp. Fig. 3C | this study |
| 200d STm$^{Aux\ \Delta hilD}$ | 200d HA902 9148 cl.2 | HA902 | Δ*asd*::tetRA Δ*dadX* Δ*alr* Δ*metC* Δ*hilD*::cat | Auxotroph Tet$^R$, Str$^R$, Cam$^R$ | Supp. Fig. 3C | this study |
| 200d STm$^{Aux\ \Delta hilD}$ | 200d HA902 9148 cl.3 | HA902 | Δ*asd*::tetRA Δ*dadX* Δ*alr* Δ*metC* Δ*hilD*::cat | Auxotroph Tet$^R$, Str$^R$, Cam$^R$ | Supp. Fig. 3C | this study |
| 200d STm$^{Aux\ \Delta hilD}$ | 200d HA902 9148 cl.4 | HA902 | Δ*asd*::tetRA Δ*dadX* Δ*alr* Δ*metC* Δ*hilD*::cat | Auxotroph Tet$^R$, Str$^R$, Cam$^R$ | Supp. Fig. 3C Supp. Fig. 3E- 200d STm$^{Aux\ \Delta hilD\ SirA\Delta 184\text{-}185}$ | this study |
| 200d STm$^{Aux\ \Delta hilD}$ | 200d HA902 9149 cl.1 | HA902 | Δ*asd*::tetRA Δ*dadX* Δ*alr* Δ*metC* Δ*hilD*::cat | Auxotroph Tet$^R$, Str$^R$, Cam$^R$ | Supp. Fig. 3C Supp. Fig. 3E- 200d STm$^{Aux\ \Delta hilD\ SirA\Delta 184\text{-}185}$ | this study |
| 200d STm$^{Aux\ \Delta hilD}$ | 200d HA902 9149 cl.2 | HA902 | Δ*asd*::tetRA Δ*dadX* Δ*alr* Δ*metC* Δ*hilD*::cat | Auxotroph Tet$^R$, Str$^R$, Cam$^R$ | Supp. Fig. 3C Supp. Fig. 3E- 200d STm$^{Aux\ \Delta hilD\ SirA\Delta 184\text{-}185}$ | this study |
| 200d STm$^{Aux\ \Delta hilD}$ | 200d HA902 9149 cl.3 | HA902 | Δ*asd*::tetRA Δ*dadX* Δ*alr* Δ*metC* Δ*hilD*::cat | Auxotroph Tet$^R$, Str$^R$, Cam$^R$ | Supp. Fig. 3C Supp. Fig. 3E- 200d STm$^{Aux\ \Delta hilD\ sirAWT}$ | this study |
| 200d STm$^{Aux\ \Delta hilD}$ | 200d HA902 9148 wp | HA902 | Δ*asd*::tetRA Δ*dadX* Δ*alr* Δ*metC* Δ*hilD*::cat | Auxotroph Tet$^R$, Str$^R$, Cam$^R$ | Supp. Fig. 3E Wp: whole population | this study |
| 200d STm$^{Aux\ \Delta hilD}$ | 200d HA902 9148 cl.1 | HA902 | Δ*asd*::tetRA Δ*dadX* Δ*alr* Δ*metC* Δ*hilD*::cat | Auxotroph Tet$^R$, Str$^R$, Cam$^R$ | Supp. Fig. 3E - 200d STm$^{Aux\ \Delta hilD\ SirA\Delta 184\text{-}185}$ | this study |
| 200d STm$^{Aux\ \Delta hilD}$ | 200d HA902 9148 cl.2 | HA902 | Δ*asd*::tetRA Δ*dadX* Δ*alr* Δ*metC* Δ*hilD*::cat | Auxotroph Tet$^R$, Str$^R$, Cam$^R$ | Supp. Fig. 3E - 200d STm$^{Aux\ \Delta hilD\ SirA\Delta 184\text{-}185}$ | this study |
| 200d STm$^{Aux\ \Delta hilD}$ | 200d HA902 9148 cl.3 | HA902 | Δ*asd*::tetRA Δ*dadX* Δ*alr* Δ*metC* Δ*hilD*::cat | Auxotroph Tet$^R$, Str$^R$, Cam$^R$ | Supp. Fig. 3E - 200d STm$^{Aux\ \Delta hilD\ SirA\Delta 184\text{-}185}$ | this study |
| 200d STm$^{Aux\ \Delta hilD}$ | 200d HA902 9148 cl.4 | HA902 | Δ*asd*::tetRA Δ*dadX* Δ*alr* Δ*metC* Δ*hilD*::cat | Auxotroph Tet$^R$, Str$^R$, Cam$^R$ | Supp. Fig. 3E- 200d STm$^{Aux\ \Delta hilD\ SirA\Delta 184\text{-}185}$ | this study |
| 200d STm$^{Aux\ \Delta hilD}$ | 200d HA902 9148 cl.5 | HA902 | Δ*asd*::tetRA Δ*dadX* Δ*alr* Δ*metC* Δ*hilD*::cat | Auxotroph Tet$^R$, Str$^R$, Cam$^R$ | Supp. Fig. 3E- 200d STm$^{Aux\ \Delta hilD\ SirA\Delta 184\text{-}185}$ | this study |
| 200d STm$^{Aux\ \Delta hilD}$ | 200d HA902 9148 cl.6 | HA902 | Δ*asd*::tetRA Δ*dadX* Δ*alr* Δ*metC* Δ*hilD*::cat | Auxotroph Tet$^R$, Str$^R$, Cam$^R$ | Supp. Fig. 3E- 200d STm$^{Aux\ \Delta hilD\ sirAWT}$ | this study |

| | | | | | | |
|---|---|---|---|---|---|---|
| 200d STm$^{Aux\ \Delta hilD}$ | 200d HA902 9148 cl.7 | HA902 | Δ*asd*::tetRA Δ*dadX* Δ*alr* Δ*metC* Δ*hilD*::cat | Auxotroph Tet$^R$, Str$^R$, Cam$^R$ | Supp. Fig. 3E- 200d STm$^{Aux\ \Delta hilD\ sirAWT}$ | this study |
| 200d STm$^{Aux\ \Delta hilD}$ | 200d HA902 9148 cl.8 | HA902 | Δ*asd*::tetRA Δ*dadX* Δ*alr* Δ*metC* Δ*hilD*::cat | Auxotroph Tet$^R$, Str$^R$, Cam$^R$ | Supp. Fig. 3E- 200d STm$^{Aux\ \Delta hilD\ sirAWT}$ | this study |
| 200d STm$^{Aux\ \Delta hilD}$ | 200d HA902 9148 cl.9 | HA902 | Δ*asd*::tetRA Δ*dadX* Δ*alr* Δ*metC* Δ*hilD*::cat | Auxotroph Tet$^R$, Str$^R$, Cam$^R$ | Supp. Fig. 3E- 200d STm$^{Aux\ \Delta hilD\ SirA\Delta 184-185}$ | this study |
| 200d STm$^{Aux\ \Delta hilD}$ | 200d HA902 9149 cl.10 | HA902 | Δ*asd*::tetRA Δ*dadX* Δ*alr* Δ*metC* Δ*hilD*::cat | Auxotroph Tet$^R$, Str$^R$, Cam$^R$ | Supp. Fig. 3E- 200d STm$^{Aux\ \Delta hilD\ SirA\Delta 184-185}$ | this study |
| 200d STm$^{Aux\ \Delta hilD}$ | 200d HA902 9149 cl.11 | HA902 | Δ*asd*::tetRA Δ*dadX* Δ*alr* Δ*metC* Δ*hilD*::cat | Auxotroph Tet$^R$, Str$^R$, Cam$^R$ | Supp. Fig. 3E- 200d STm$^{Aux\ \Delta hilD\ SirA\Delta 184-185}$ | this study |
| 200d STm$^{Aux\ \Delta hilD}$ | 200d HA902 9149 cl.12 | HA902 | Δ*asd*::tetRA Δ*dadX* Δ*alr* Δ*metC* Δ*hilD*::cat | Auxotroph Tet$^R$, Str$^R$, Cam$^R$ | Supp. Fig. 3E- 200d STm$^{Aux\ \Delta hilD\ sirAWT}$ | this study |
| 200d STm$^{Aux\ \Delta hilD}$ | 200d HA902 9149 cl.13 | HA902 | Δ*asd*::tetRA Δ*dadX* Δ*alr* Δ*metC* Δ*hilD*::cat | Auxotroph Tet$^R$, Str$^R$, Cam$^R$ | Supp. Fig. 3E- 200d STm$^{Aux\ \Delta hilD\ SirA\Delta 184-185}$ | this study |
| 200d STm$^{Aux\ \Delta hilD}$ | 200d HA902 9149 cl.14 | HA902 | Δ*asd*::tetRA Δ*dadX* Δ*alr* Δ*metC* Δ*hilD*::cat | Auxotroph Tet$^R$, Str$^R$, Cam$^R$ | Supp. Fig. 3E- 200d STm$^{Aux\ \Delta hilD\ sirAWT}$ | this study |
| 200d STm$^{Aux\ \Delta hilD}$ | 200d HA902 9149 cl.15 | HA902 | Δ*asd*::tetRA Δ*dadX* Δ*alr* Δ*metC* Δ*hilD*::cat | Auxotroph Tet$^R$, Str$^R$, Cam$^R$ | Supp. Fig. 3E- 200d STm$^{Aux\ \Delta hilD\ sirAWT}$ | this study |
| 200d STm$^{Aux\ \Delta hilD}$ | 200d HA902 9149 cl.16 | HA902 | Δ*asd*::tetRA Δ*dadX* Δ*alr* Δ*metC* Δ*hilD*::cat | Auxotroph Tet$^R$, Str$^R$, Cam$^R$ | Supp. Fig. 3E- 200d STm$^{Aux\ \Delta hilD\ sirAWT}$ | this study |
| 1461d STm$^{Aux}$ | 1461d STm$^{Aux}$ | HA630 cl.3 | Δ*asd*::tetRA Δ*dadX* Δ*alr* Δ*metC* | Auxotroph Tet$^R$, Str$^R$ | Fig. 4A | this study |
| 1461d STm$^{Aux}$ | 1461d STm$^{Aux}$ 1K | HA630 cl.3 | Δ*asd*::tetRA Δ*dadX* Δ*alr* Δ*metC* | Auxotroph Tet$^R$, Str$^R$, Kan$^R$ | Fig. 2 | this study |
| 1461d STm$^{Ex\text{-}Aux}$ | HA755 | 1461d STm$^{Aux}$ | Δ*dadX* Δ*metC* | Ex-Auxotroph Str$^R$ | Fig. 4A, B Fig. 6 | this study |
| 1461d STm$^{Ex\text{-}Aux\ \Delta ssaV}$ | 1461d STm$^{Ex\text{-}Aux\ \Delta ssaV}$ | HA755 | Δ*dadX* Δ*metC* Δ*ssav*::cat | Ex-Auxotroph Str$^R$, Cam$^R$ | Fig. 5 | this study |
| 1842d STm$^{Aux}$ | 1842d 433 cl.1 | HA630 cl.3 | Δ*asd*::tetRA Δ*dadX* Δ*alr* Δ*metC* | Auxotroph Tet$^R$, Str$^R$ | Fig. 4A- STm$^{Aux\ SirA\Delta 184-185}$ | this study |
| 1842d STm$^{Aux}$ | 1842d 433 cl.2 | HA630 cl.3 | Δ*asd*::tetRA Δ*dadX* Δ*alr* Δ*metC* | Auxotroph Tet$^R$, Str$^R$ | Fig. 4A- STm$^{Aux\ SirA\Delta 184-185}$ | this study |
| 1842d STm$^{Aux}$ | 1842d 433 cl.3 | HA630 cl.3 | Δ*asd*::tetRA Δ*dadX* Δ*alr* Δ*metC* | Auxotroph Tet$^R$, Str$^R$ | Fig. 4A- STm$^{Aux\ SirA\Delta 184-185}$ | this study |
| 1842d STm$^{Aux}$ | 1842d 515 cl.1 | HA630 cl.3 | Δ*asd*::tetRA Δ*dadX* Δ*alr* Δ*metC* | Auxotroph Tet$^R$, Str$^R$ | Fig. 4A- STm$^{Aux\ SirA\Delta 184-185}$ | this study |

| | | | | | | |
|---|---|---|---|---|---|---|
| 1842d $STm^{Aux}$ | 1842d 515 cl.2 | HA630 cl.3 | Δ*asd*::tetRA Δ*dadX* Δ*alr* Δ*metC* | Auxotroph $Tet^R$, $Str^R$ | Fig. 4A- $STm^{Aux}_{SirA\Delta 184-185}$ | this study |
| 1842d $STm^{Aux}$ | 1842d 520 cl.1 | HA630 cl.3 | Δ*asd*::tetRA Δ*dadX* Δ*alr* Δ*metC* | Auxotroph $Tet^R$, $Str^R$ | Fig. 4A - $STm^{Aux}_{SirA\Delta 184-185}$ | this study |
| 1842d $STm^{Ex\text{-}Aux}$ | 1842d 520 cl.$1^{Ex\text{-}Aux}$ | HA630 cl.3 | Δ*dadX* Δ*metC* | Ex-Auxotroph $Str^R$ | Fig. 4A, B<br>Fig. 6 | this study |
| 1842d $STm^{Aux}$ | 1842d 520 cl.2 | HA630 cl.3 | Δ*asd*::tetRA Δ*dadX* Δ*alr* Δ*metC* | Auxotroph $Tet^R$, $Str^R$ | Fig. 4A - $STm^{Aux}_{SirA\Delta 184-185}$ | this study |
| 1842d $STm^{Aux}$ | 1842d 520 cl.3 | HA630 cl.3 | Δ*asd*::tetRA Δ*dadX* Δ*alr* Δ*metC* | Auxotroph $Tet^R$, $Str^R$ | Fig. 4A - $STm^{Aux}_{SirA\Delta 184-185}$ | this study |
